# Intricate genetic programs controlling dormancy in *Mycobacterium tuberculosis*

**DOI:** 10.1101/709378

**Authors:** Abrar A. Abidi, Eliza J. R. Peterson, Mario L. Arrieta-Ortiz, Boris Aguilar, James T. Yurkovich, Amardeep Kaur, Min Pan, Vivek Srinivas, Ilya Shmulevich, Nitin S. Baliga

**Affiliations:** Institute for Systems Biology, 401 Terry Avenue North, Seattle, Washington 98109, USA

## Abstract

*Mycobacterium tuberculosis* (MTB), responsible for the deadliest infectious disease worldwide, displays the remarkable ability to transition in and out of dormancy, a hallmark of the pathogen’s capacity to evade the immune system and opportunistically exploit immunocompromised individuals. Uncovering the gene regulatory programs that underlie the dramatic phenotypic shifts in MTB during disease latency and reactivation has posed an extraordinary challenge. We developed a novel experimental system to precisely control dissolved oxygen levels in MTB cultures in order to capture the chain of transcriptional events that unfold as MTB transitions into and out of hypoxia-induced dormancy. Using a comprehensive genome-wide transcription factor binding location map and insights from network topology analysis, we identified regulatory circuits that deterministically drive sequential transitions across six transcriptionally and functionally distinct states encompassing more than three-fifths of the MTB genome. The architecture of the genetic programs explains the transcriptional dynamics underlying synchronous entry of cells into a dormant state that is primed to infect the host upon encountering favorable conditions.

**One Sentence Summary:** High-resolution transcriptional time-course reveals six-state genetic program that enables MTB to enter and exit hypoxia-induced dormancy.

## Main Text

*Mycobacterium tuberculosis* (MTB) kills more people than any other infectious agent, causing ∼10 million new cases of active tuberculosis (TB) disease and 1.7 million deaths each year (Murray et al., 2014). TB remains a major human public health burden, in large part due to the sizeable reservoir of latently infected individuals, who may relapse into active disease decades after acquiring the infection. MTB can persist in a stable, non-replicative (often termed dormant) state within the host for months or years without symptoms, and then revive to initiate the production of lesions and active TB disease. Moreover, dormant cells may be responsible for the slow treatment response of patients with active TB. Elucidation of the factors that affect treatment outcome, latency and activation requires a better characterization of functional states adopted by the pathogen during progression of the disease, as well as a mechanistic understanding of the genetic programs that orchestrate transitions between these states.

Hypoxia, an environmental stress encountered by MTB within granulomas (Tsai et al., 2006), is sufficient to shift the pathogen into a defined non-growing survival form, which can be reversed upon aeration of the culture (Chao and Rubin, 2010). Therefore, hypoxia has been leveraged as an *in vitro* approximation to study MTB dormancy and the underlying genetic programs. However, previous transcriptional analyses under *in vitro* hypoxic conditions (via the Wayne model in which MTB cultures are sealed and gradually depleted of oxygen (Wayne and Hayes, 1996; Wayne and Sohaskey, 2001) or the defined hypoxia model in which nitrogen gas is flowed into the headspace to rapidly deplete oxygen (Kempner, 1939; Yuan et al., 1998)) were limited to either static snapshots, or low-resolution time-course studies (Muttucumaru et al., 2004; Rustad et al., 2008; Sherman et al., 2001). Moreover, deletion of previously identified transcriptional regulators thought crucial to hypoxia-induced dormancy (i.e. Δ*dosR*, Δ*sigE*Δ*sigH*), conferred only mild growth defects under hypoxic conditions (Boon and Dick, 2002; Rustad et al., 2008; Rustad et al., 2009), suggesting a genetic circuit architecture that has evolved to withstand environmental and genetic perturbations. Here, we developed a novel experimental platform to characterize MTB’s response to changing oxygen (O_2_) levels in considerably more depth. We reveal detailed transcriptional dynamics and coordinated regulatory circuits that enable the pathogen’s transition into and out of hypoxia-induced dormancy.

To detail the genetic programs underlying hypoxia-induced dormancy in MTB, we needed to obtain accurate dynamic measurements of genome-wide expression, over an O_2_ gradient. Previous experimental models to study O_2_ tension and growth arrest in MTB were not suitable for the accuracy and resolution of measurements needed. In particular, the Wayne model has issues with reproducibility (Rustad et al., 2008) and the defined hypoxia model depletes O_2_ very quickly, thereby hindering high-resolution sampling during critical transition periods. Moreover, neither model has been performed with real-time monitoring of O_2_ levels to accurately relate the transcriptional state of MTB with a precise O_2_ measurement. Therefore, we designed a new programmable multiplexed reactor system, the controlled O_2_ model, to precisely manipulate and monitor O_2_ levels within the growth medium–even during sampling (**Fig 1A**). The precise control engineered into the system enabled high-resolution sampling across a time-course and O_2_ gradient, with minimal disturbance to the bacteria and high reproducibility across culture replicates and experiments (**Fig S1**). Briefly, air and nitrogen (N_2_) gas lines were connected to separate mass flow controllers, which allowed for programmable gradients of gas mixtures to be streamed into the headspace of spinner flasks containing MTB in media. Moreover, we used O_2_ sensor spots and fiber optic technology to non-invasively measure the dissolved O_2_ content of the cultures. Both the mass flow controllers and O_2_ sensor spots were configured for remote management, advantageous for a biosafety level 3 pathogen. With the controlled O_2_ model, we performed a time-course experiment, which involved a steady depletion of dissolved oxygen (DO) over 2 days from full aeration (∼80% DO) to hypoxia (0% DO). This steady depletion was achieved by programming the mass flow controllers to produce the desired mixture of air and N_2_. The cultures were maintained in hypoxia for 2 days by streaming only N_2_, then reaerated over 1 day by a programmed increase in air flow (**Fig 1B**). Over the time-course, we harvested samples in triplicates, one each from three independent reactors, via sampling ports that prevented aeration of the culture. We sampled at high frequency during the period when cultures transitioned from 10% to 0% DO, as well as from 0% to 10% DO and at lower but regular frequency across the remaining 120-hour experiment. Samples were flash frozen in liquid N_2_ and later processed for gene expression profiling by RNA-sequencing.

**Figure 1.**
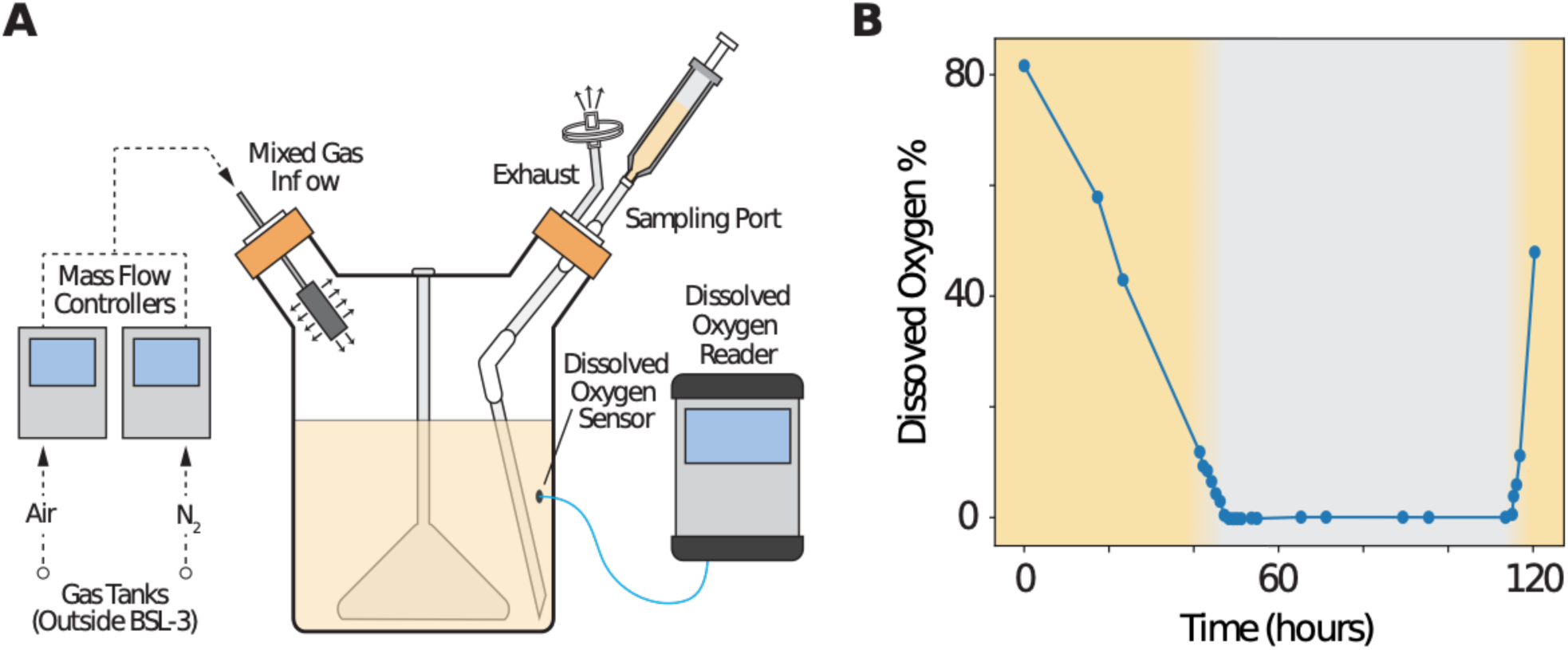
Schematic of the controlled O_2_ model reactor system and DO profiles. **(A)** Programmable mass flow controllers were used to modulate the ratio of air and nitrogen (N_2_) in a gas mixture that was flowed into the headspace of spinner flasks containing cultures of MTB. Dissolved oxygen sensor spots and fiber optic technology non-invasively provided real time and remote readout of the dissolved oxygen (DO) levels within the culture media. Samples were drawn from a sampling port attached to one of the side arms of the spinner flask. Four reactors were multiplexed and individually monitored for DO levels to obtain biological replicates. **(B)** DO levels across the 120-hour time course. Points are the average of three biological replicates; the yellow shading indicate the periods of controlled O_2_ depletion and reaeration, whereas the white background indicates a sustained 2 day immersion in hypoxia.

Over the course of the experiment, nearly 64% of all genes (non-coding regions were excluded) in the MTB genome were significantly differentially expressed (2,582 genes with adjusted *P*-value < 0.05 and estimated absolute log2 fold-change >1). The number of differentially expressed genes from our controlled O_2_ system is an order of magnitude greater than the number from earlier microarray studies using the Wayne (Muttucumaru et al., 2004) (299 genes) or defined hypoxia (Rustad et al., 2008) (274 genes) models. Nevertheless, there is significant overlap across gene sets between the three models of hypoxia-induced dormancy (**Table S1**). Interestingly, the controlled O_2_ model significantly recapitulated differential expression observed from intracellular MTB (Peterson et al., 2019) (enrichment test *P*-value=4.29 × 10^−30^), whereas the other hypoxia models did not or had a low recall of the differentially expressed genes (**Table S2**). These findings highlight the capability of our model to capture MTB’s transcriptional programs during dormancy that are relevant to MTB within host cells.

To further characterize the MTB transcriptional states over the time course and O_2_ gradient, we applied dimensionality reduction techniques that allowed us to define tightly clustered samples (**Fig S2;** *see methods*). The six identified clusters are shown in a two-dimension t-distributed stochastic neighbor embedding (tSNE) plot (**Fig 2A**). Each cluster represents a distinct transcriptional state and was associated with sets of non-overlapping differentially expressed genes: Normoxia (81 genes), Depletion (446 genes), Early hypoxia (328 genes), Mid hypoxia (320 genes), Late hypoxia (978 genes), and Resuscitation (429 genes) (**Dataset S1**). Each differentially expressed gene was assigned to the state in which it had the highest mean expression (*see methods*). The average expression profiles for the gene sets reveal that the states transition from one to another and that transitions are oxygen-(e.g. Late hypoxia into Resuscitation occurred upon re-introducing air into the culture) and time-dependent (e.g. Early/Mid hypoxia into Late hypoxia occurred ∼40 h after the culture reached 0% DO) (**Fig 2B**). As such, the six states were also defined with oxygen and time intervals (**Table S3**), with the exception of 46-49 h, where there was oscillation between Early hypoxia and Late hypoxia states as the culture went below ∼3% DO (**Fig 2**). While this intriguing “flicker” behavior could be experimental noise, these anomalous time points (measured roughly one hour apart) clearly cluster with Late hypoxia. Such oscillatory expression could be generated by inherent properties of the network structure, which we describe later.

**Figure. 2.**
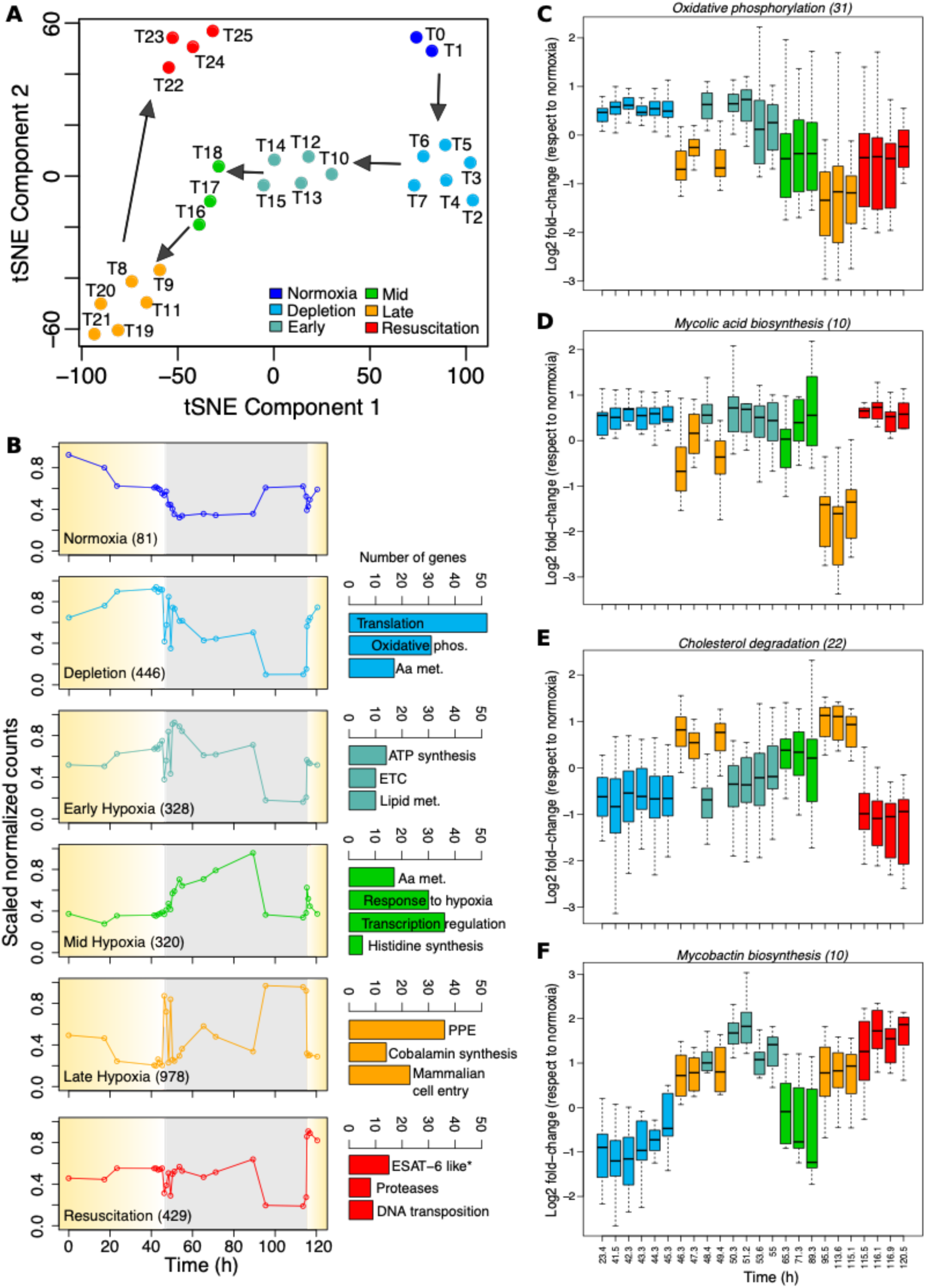
The controlled O_2_ model captures distinct cell states over time course and O_2_ gradient. **(A)** t-SNE analysis of all samples across the time course and hypoxia gradient. **(B)** Average expression profiles for state-specific gene sets across the time course and hypoxia gradient. The yellow shading indicate the periods of controlled O_2_ depletion and reaeration, whereas the grey background indicates a sustained 2 day immersion in hypoxia. General theme of significant functional term clusters defined by DAVID (Huang da et al., 2009), in each state are indicated. The star symbol (*) indicates the most enriched term at the individual term level in the Resuscitation state (not present in any of the significant term clusters). (**C-F**) Log2 fold-change respect to normoxia (T0 and T1) of selected metabolic pathways reconstructed by Kavvas et al (Kavvas et al., 2018); numbers in parentheses indicated the number of genes in the relevant pathway.

Genes associated to the Depletion state (DO between 43% and 4%) were enriched for growth-related functions including amino acid metabolism, oxidative phosphorylation and translation (**Fig 2B**). In Early hypoxia, ATP synthase and genes involved in electron transport chain and lipid metabolism were highly enriched and expressed, even more so than in Normoxia. Furthermore, these metabolic genes were then significantly down regulated during Late hypoxia. This result indicates that Early hypoxia is a metabolically active state that may exist for MTB to prepare itself for an upcoming metabolically quiescent state (i.e. Late hypoxia). Mid hypoxia genes were enriched in stress response genes, indicating the bacteria are sensing and adapting to the anaerobic environment. In Late hypoxia, genes essential for MTB to infiltrate host cells were induced. Furthermore, genes for 32 proteins that belong to the proline-glutamic acid (PE) and proline-proline-glutamic acid (PPE) family, whose functions remain largely unknown (Bottai and Brosch, 2009) were up-regulated. These PPE family proteins have been proposed to modulate the host’s immune response (Tiwari et al., 2012), generate antigenic variation (Cole et al., 1998) and were shown to be secreted by MTB’s ESX-5 export system (Abdallah et al., 2008). Interestingly, genes encoding the components of ESX-5 export system (as well as ESX-1 and ESX-3) were rapidly activated as soon as MTB shifted from Late hypoxia into Resuscitation, minutes after air was introduced back into the culture (**Fig S3**). It is possible that Late hypoxia not only engenders quiescence in MTB but also sequesters a collection of PPE proteins in anticipation of resuscitation and ESX system production. In the Resuscitation state, proteases, transposases and insertion sequences were also enriched among activated genes. These functional groups suggest that MTB may strategically avert immune recognition through antigenic heterogeneity (via ESX secretion of PPE proteins) and simultaneously reorganize its genome (via transposases and insertion sequences) to increase its chances for survival and transmission to a new host upon resuscitation.

Using the gene expression data, we also analyzed changes in MTB metabolic pathways, as reconstructed by Kavvas et al (Kavvas et al., 2018), along the hypoxia time series. The expression of many metabolic pathways reiterated the state transitions described above (**Fig 2C-F**). For example, significant down-regulation of genes involved in oxidative phosphorylation during Late hypoxia (**Fig 2C**), confirming the loss of energy related pathways during Late hypoxia. Furthermore, MTB’s dependency on alternative carbon sources was also observed in Late hypoxia, with significant up-regulation of genes related to cholesterol degradation and a simultaneous down-regulation of mycolic acid biosynthesis genes (**Fig 2D-E**). Reaeration of the MTB culture and entry into the Resuscitation stage reversed these expression trends of late hypoxia. Additionally, we observed interesting expression dynamics for genes involved in mycobactin biosynthesis, where expression was generally up-regulated during hypoxia states and Resuscitation, excluding Mid hypoxia (**Fig 2F**). The increase in mycobactin, an iron chelator, is essential for MTB to access iron, particularly when the bacteria competes with the host for the metal (McMahon et al., 2012). Overall, MTB adaptation to hypoxia involves rewiring of several metabolic pathways, indicating an evolutionarily learned and coordinated response to stresses that typically co-exist within the host environment (e.g., hypoxia, starvation, and iron limitation), despite the singular *in vitro* perturbation.

The six-state model across the time course and O_2_ gradient revealed distinct patterns of expression suggestive of intriguing and coordinated regulatory programs. Several methods are available for reconstructing gene regulatory networks (GRN) along time series expression data (Baugh et al., 2005; Bromberg et al., 2008; Luscombe et al., 2004). We selected DREM 2.0 (Schulz et al., 2012), which has been successfully applied to various systems (e.g., fly (Consortium et al., 2010), yeast (Ernst et al., 2007), *E. coli* (Ernst et al., 2008)) and is ideal for identifying dynamic transcriptional events over time and perturbations. The Dynamic Regulatory Events Miner (DREM) integrates time series and snapshots of the GRN of interest using an input-output Hidden Markov Model (Ernst et al., 2007). In so doing, DREM learns a dynamic GRN by identifying bifurcation points—places in the time series where a group of co-expressed genes begins to diverge. These bifurcation points are annotated with the proposed TFs controlling the split, leading to a combined dynamic model. Using the hypoxia time course expression dataset and a TF-target gene network derived from the ChIP-seq assessment of 154 TFs overexpressed in MTB (Minch et al., 2015), DREM identified bifurcation points that coincide with transitions between the six states (**Fig 3A** and **Fig S4**). The bifurcation points defined by DREM reinforce the importance of transcriptional regulation in the progression between states. DREM identified TFs that are known to mediate MTB’s response to hypoxia (e.g., DosR, Rv0081, Rv0324) (Galagan et al., 2013) along with additional TFs with a potential role in hypoxia. In particular, Rv1353c stood out for being the only TF linked to the time points that precede and mark the end of Late hypoxia. The Rv1353c regulon is the third largest, with 596 genes (after the Rv0081 and Rv0678 regulons), and DREM predicts that Rv1353c regulatory activity may be important for sustained hypoxic conditions. In addition, DREM suggests that CsoR (Rv0967), a TF that controls MTB’s response to copper stress (Marcus et al., 2016), may also have an unappreciated role in hypoxia. Interestingly, DREM associated CsoR with the bifurcation points preceding Early hypoxia and Resuscitation. In the latter point, *csoR* transcriptional level transitioned from its lowest value (Late hypoxia) to its highest value (Resuscitation). This suggests a potential bifunctional activity of CsoR in controlling MTB’s transcriptional response both in and out of hypoxia. In fact, 87% of DEGs (223 genes in total) from a *csoR* knockout mutant (Marcus et al., 2016) were differentially expressed at some point during the time course and oxygen gradient (*P*-value = 1.7 × 10^−16^). Specifically, this set of DEGs was enriched with members of Early hypoxia, Mid hypoxia, and Resuscitation, supporting the CsoR bifurcation points predicted by DREM. We propose further investigation to evaluate the consequences of perturbing Rv1353c and CsoR activity during hypoxia adaptation.

**Fig. 3.**
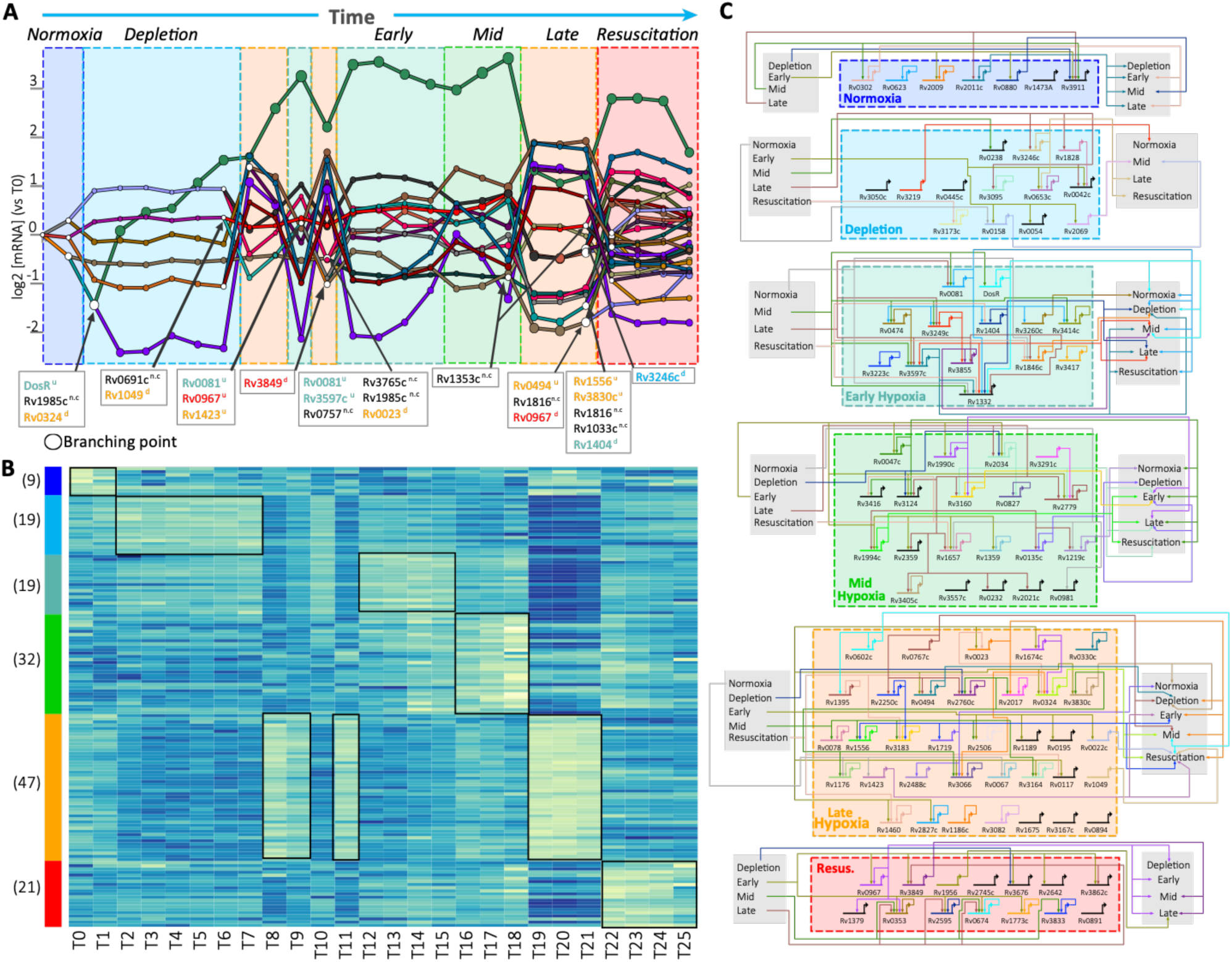
Transcriptional circuits controlling the six transcriptional states adopted by MTB during entry and exit from hypoxia. (**A**) DREM output. Transcription factors (TFs) associated with selected branching points (white nodes) are shown. Time points and TFs are colored code based on transcriptional states membership. The *n.c, d* and *u* superscripts indicate no change, down-regulation and up-regulation, respectively. (**B**) Heatmap with transcriptional profiles of 147 differentially expressed MTB TFs; number in parentheses indicate the number of TFs associated with each state. (**C**) TF-TF network of differentially expressed TFs in the controlled O_2_ model. ChIP-seq derived protein-DNA interactions reported in Minch *et a*l (Minch et al., 2015) were used to establish the connections between TFs. Only TFs with one or more differentially expressed targets were included in the diagram. The diagram was generated with the Biotapestry tool (Paquette, 2016).

While DREM was able to identify key TFs involved in state transitions with >0% DO, only a single bifurcation point (associated to Rv1353c) was predicted within the hypoxia window between T12 and T20. This single TF prediction during hypoxic conditions reveals a limitation of DREM reconstructions, which is the focus on bifurcation points. Although there are clear changes in expression in sets of co-expressed genes during these hypoxic stages (e.g., purple path in **Fig 3A**), DREM does not associate a TF with these changes due to the absence of a bifurcation in the gene set. Instead, the genes continue to change expression as a unit and could be influenced by a TF whose activity is being modified over time in hypoxia. Another explanation for the lack of TFs identified by DREM during the hypoxia-associated states (Early, Mid, Late hypoxia) could be the normoxic conditions used for collecting the available protein-DNA interaction data (Minch et al., 2015).

The set of TFs identified by DREM included only 16% of the 147 putative TFs differentially expressed at some point across the time course and oxygen gradient (**Fig 3B**). In fact, Late hypoxia contains 24% of all differentially expressed TFs (47 TFs). The large number of differentially expressed TFs suggested complex and combinatorial circuitry patterns could be involved in MTB’s adaptation to hypoxic conditions. **Fig 3C** shows the dense TF-TF connectivity within and between transcriptional states, according to available protein-DNA binding data (Minch et al., 2015). The key regulatory proteins of the TF-TF network were identified based on betweenness centrality, which characterizes the connectivity of interacting nodes in the network. The top high-degree nodes were (in decreasing order) Rv0081 (Early hypoxia), Rv3597c (Lsr2; Early hypoxia), Rv1990c (Mid hypoxia), Rv2034 (Mid hypoxia) and Rv0023 (Late hypoxia). As high-degree nodes, drugs targeting one or more of these regulatory hubs may have a major impact on MTB survival. In support of this, Bartek and collaborators showed that deletion of *lsr2* significantly compromised adaptation of MTB to hypoxic conditions (Bartek et al., 2014). Notably, Lsr2 had the second and third largest outdegree (number of TF targets) and indegree (number of transcriptional regulators), respectively. Lsr2 directly controls TFs from Depletion (one TF), Early hypoxia (two TFs), Mid hypoxia (four TFs), Late hypoxia (five TFs) and Resuscitation (one TF). Moreover, *lsr2* is regulated by Mid hypoxia TFs (Rv1994c, Rv2034 and Rv3160c) and Late hypoxia TFs (Rv0023, Rv0324 and Rv1460). The critical role of Lsr2 in the coordination between hypoxia-related states offers an explanation for the known importance of Lsr2 in hypoxic conditions.

The high connectivity of the TF-TF network revealed regulatory hubs that activate one state while repressing another. Interestingly, DREM also identified bifurcation points in DO >0% with down-regulated Late hypoxia TFs (Rv0023c, Rv0324 and Rv1049), indicating a concurrent repression of Late hypoxia regulators and activation of earlier states. Motivated by these findings, we evaluated the enrichment of regulons of all differentially expressed TFs with members of each transcriptional state. Many regulons (n = 21) were significantly enriched with members of their TF’s state (permutation test *P*-value = 0), while even more regulons (n = 49, permutation test *P*-value = 0) were significantly enriched with members of other transcriptional states (**Dataset S2**). In other words, TFs act to express members of their own state while repressing members of another state (shown in TF-TF interactions of **Fig 3C**). Such behavior by TFs, described as mutual inhibition (Gardner et al., 2000; Glass and Kauffman, 1973; Huang et al., 2007), proposes a mechanism for the coordination and cooperation between transcriptional states to achieve the proper timing and gene expression levels to successfully adapt to changes in O_2_.

Interactions among TFs can also form specific network motifs that perform defined dynamical functions in response to changing environmental conditions (Alon, 2007; Shiraishi et al., 2010; Wu et al., 2011). Network motifs, such as feedforward loops (FFLs), single-input modules, or bistable toggle switches (Alon, 2007) are recurring gene patterns found within gene regulatory networks. To unbiasedly search for network motifs that may be involved in hypoxia adaptation, we analyzed the experimentally determined MTB TF-target gene interactions from ChIP-seq (Minch et al., 2015) using FANMOD (Wernicke and Rasche, 2006) in the MotifNet Webserver (Smoly et al., 2017). We found the MTB ChIP-seq network is significantly enriched with FFLs, a common network motif composed of two input TFs, one of which regulates the other and both of which jointly regulate a target gene (or set of genes) (Mangan and Alon, 2003). We ran extensive permutation tests to confirm the likelihood (*P*-value = 0.001) of 1690 FFL instances emerging in a random network with the same number of nodes and edges. Interestingly, Rv0081 is the most frequent regulator at the “top” of the FFLs (45.4% of all detected instances) and also has the highest-degree connectivity (described above). Rv0081 has been previously linked to MTB’s response to hypoxia (Galagan et al., 2013; Prosser et al., 2017) and is itself a target gene of the well-characterized regulator of dormancy survival, DosR (Boon and Dick, 2002; Park et al., 2003; Sherman et al., 2001; Vasudeva-Rao and McDonough, 2008). To evaluate the involvement of Rv0081-centered FFLs in the transcriptional changes observed during hypoxia, we explored some of the most frequent TF pairs found in FFL configuration). The top pair, Rv0081-Rv0324 controls 134 genes significantly enriched with Late hypoxia genes (*P*-value = 6.4×10^−5^). Rv0081 also frequently pairs with Rv3249 and controls 87 genes enriched with Depletion state genes (*P*-value = 1.1×10^−5^). Another frequent pair combines Rv0023 and Rv0324 to control 70 genes enriched in Mid hypoxia genes (*P*-value = 3×10^−5^). We explored the directionality of these state-specific FFL target genes using gene expression data from MTB TF overexpression (TFOE) strains in normoxia (Rustad et al., 2014) and a MTB Rv0081 gene deletion (ΔRv0081) strain in hypoxia (Sun et al., 2018). For example, the majority of Depletion genes with differential expression in the ΔRv0081 strain were up-regulated, suggesting a negative relationship with Rv0081 in hypoxia (**Fig 4A**). Moreover, we observed that Depletion genes controlled by the Rv0081-Rv3249c FFL were significantly down-regulated in the Rv0081 TFOE strain (**Fig 4B**). In contrast, there is a positive relationship between Rv0081and Late hypoxia genes during hypoxia as indicated by a largely decreased expression of Late hypoxia genes in the ΔRv0081 strain (**Fig 4C**). Furthermore, Late hypoxia genes controlled by the Rv0081-Rv0324 FFL were significantly up-regulated (**Fig 4D**). Altogether, we generated a model of interlocking FFLs that together up-regulate 213 genes corresponding to Late hypoxia, while also coordinating the repression of Mid hypoxia and Depletion genes (**Fig 4E**). The overlapping sets of network motifs act to reinforce each other’s function and direct the complex physiological state transitions required to adapt to decreasing DO levels.

**Fig. 4.**
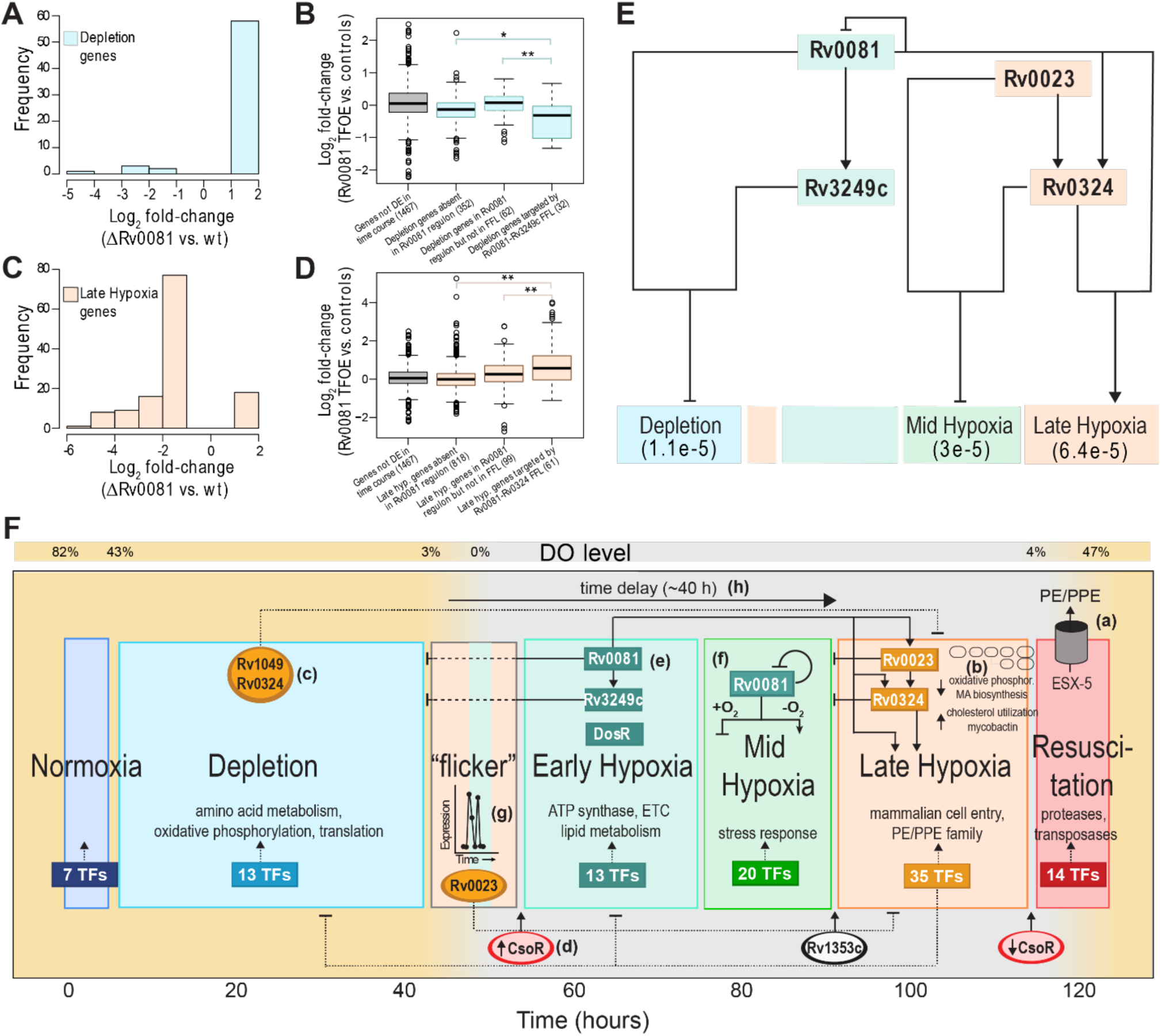
Data supporting Rv0081-controlled interlocking FFLs and overview of the transcriptional dynamics across the six-state model that enables MTB to enter and exit hypoxia-induced dormancy. (**A**) The log2 fold-change distribution of Depletion genes with significant differential expression (*P*-value < 0.05, absolute log2 fold-change >1) from the ΔRv0081 strain in hypoxia (Sun et al., 2018). (**B**) Boxplots representing log2 fold-change of various gene groups related to Depletion state from Rv0081 transcription factor overexpression (TFOE) (Rustad et al., 2014). The number in parentheses indicates the number of genes evaluated in each group. (**C**) The log2 fold-change distribution of Late hypoxia genes with significant differential expression (*P*-value < 0.05, absolute log2 fold-change >1) from the ΔRv0081 strain in hypoxia (Sun et al., 2018). (**D**) Boxplots representing log2 fold-change of various gene groups related to Late hypoxia state from Rv0081 transcription factor overexpression (TFOE) (Rustad et al., 2014). (**E**) Model of Rv0081-controlled interlocking FFLs that together up-regulate a significant number of genes corresponding to Late hypoxia, while also repressing Mid hypoxia and Depletion genes. The *P*-values for enrichment of the FFL-controlled genes from each state are indicated below the state name, as evaluated using a hypergeometric test. (**F**) Summary overview of the transcriptional dynamics, inferred key regulators, and regulatory circuits that were revealed from the high-resolution and longitudinal gene expression profiling across the O_2_ gradient. Key features described in the text include: (a) ESX-5 secretion of PE/PPE proteins that were expressed during Late hypoxia; (b) changes in metabolic pathways during Late hypoxia; (c) DREM predicted TFs (shown as ovals throughout figure) that concurrently repress Late hypoxia genes and activate earlier states (mutual inhibition connections shown as dotted lines throughout figure); (d) DREM identified CsoR in controlling MTB’s transcriptional response both in and out of hypoxia; (e) Rv0081-controlled interlocking FFLs; (f) the regulatory activity of Rv0081 is oxygen-dependent; (g) the “flicker” of Late hypoxia gene expression as DO dropped below 3%; (h) the shift to Late hypoxia occurred after 40 hours in hypoxia. * indicates *P*-value < 0.05, ** indicates *P*-value < 0.01

Three important hypotheses developed from identifying network motif topology. The first hypothesis is that Rv0081 plays a pivotal role in the adaptation of MTB to hypoxia (Galagan et al., 2013) and its regulatory activity may be oxygen-dependent. The bifunctional activity of Rv0081 is based on comparison between the normoxic TFOE data (Rustad et al., 2014) and hypoxic ΔRv0081 data (Sun et al., 2018) (**Fig S5**). The regulatory targets of Rv0081 had very little concordance of fold-change expression in these different conditions. Most genes that were significantly down-regulated in the ΔRv0081 strain in hypoxia showed no fold-change difference in the TFOE data (in normoxia). This observation supports recent work demonstrating that Rv0081 had altered DNA-binding ability under hypoxic conditions, with evidence that formate ion accumulation and/or post-translational modifications may be involved in the conditional regulatory activity (Kumar et al., 2019). Furthermore, dual activity of Rv0081 seems to be necessary for coordinating the repression of Late hypoxia TFs in early hypoxic time points and their up-regulation in Late hypoxia (**Fig S6**).

The second hypothesis is that Rv0081 may be involved in the state oscillations observed as the DO dropped below 3% (**Fig 2**). Under these low O_2_ conditions, the MTB transcriptome oscillated between two states: Late hypoxia genes were expressed (T8 and T9), then Early hypoxia genes (T10), then back to Late hypoxia (T11), before ultimately committing to Early hypoxia (T12-T18). This “flicker” between Early hypoxia and Late hypoxia, measured roughly one hour apart, could emerge from oscillatory mechanisms involving an Rv0081-directed incoherent FFL (I-FFL) (Geva-Zatorsky et al., 2006; Kholodenko, 2000; Novak and Tyson, 2008). In I-FFLs, one TF acts positively while the other TF acts negatively, resulting in a pulse of target gene(s) expression. Interestingly, the peak height between the first and second pulse of Late hypoxia genes had roughly equal normalized expression (**Fig 2B & Fig S7)**, suggesting a potential detection of fold-change based on the I-FFL, as described by Goentoro and colleagues (Goentoro et al., 2009). Fitting Rv0081 and Early Hypoxia TFs (excluding Rv0081) between T7-T20 with three configurations of I-FFLs (cooperative, independent and exclusive; *see methods*), cooperative TF binding most closely modeled the observed average expression of Late hypoxia genes (**Fig S7A**, RMSD = 0.0965). Importantly, all three I-FFL configurations were able to reproduce the oscillatory expression of Late hypoxia genes when DO dropped below 3%, concluding that the I-FFL motif can explain the observed “flicker” between Early and Late hypoxia states. Furthermore, the parameters of the exclusive binding model are in the fold change detection region (**Fig S7C**), indicating that a fold-change detection mechanism is possible if a subset of repressors interact exclusively with Rv0081 to regulate some Late hypoxia genes. While the role of the I-FFL as a fold-change detector requires further exploration, it is intriguing to hypothesize that by sensing relative changes in gene expression nearing hypoxia, a potentially variable and “noisy” period, the circuit could serve to synchronize the hypoxic response across all cells in the population.

Finally, the third hypothesis is that the I-FFL controlled by Rv0081 also regulates the transition to Late hypoxia. Late hypoxia accounts for the largest change in expression and the transcriptional state most characteristic of dormant (i.e. nonreplicating) MTB (Schnappinger et al., 2003; Voskuil et al., 2003). In addition to reproducing the “flicker”, all three I-FFL configurations modeled a delay in Late hypoxia gene expression after entering hypoxia **(Fig S7**). The time scale of the delay element, about 40 hours from entering 0% DO to Late hypoxia transition, is consistent with delayed translation observed in slow-growing MTB in response to nitric oxide (Cortes et al., 2017). It was recently demonstrated that delayed regulatory interactions within I-FFLs (with mutual inhibition present) produced state transitions related to T cell exhaustion, after a fixed time post-stimulation (Bolouri, 2019). As such, the delay element may function in MTB to incorporate robustness into the hypoxic response, ensuring that Late hypoxia (with large-scale expression changes) is not activated prematurely. Further investigation is required to determine whether the duration of the delay element is fixed or variable in a manner dependent on how MTB enters hypoxia. The shift to Late hypoxia, after 40 hours in hypoxia and following transition through two intermediate hypoxic states, is one of the most intriguing revelations from this study and required the development of a novel reactor system and the high-resolution profiling that was performed here. The elucidation of regulatory circuits that control the large, altered transcriptome of Late hypoxia offers novel drug targets that could block the underlying mechanisms that contribute to replication suppression, alternate respiratory/metabolic pathways, and phenotypic tolerance associated with dormant MTB.

This report presents the high-resolution system-wide gene expression profiling of MTB across a 5-day time course of hypoxia and reaeration. A novel reactor system was designed to allow for exquisite control and monitoring of O_2_ levels, thereby uncovering intermediate transcriptional states and dynamic expression patterns, not previously described. Gene expression profiling revealed that three-fifths of all genes in the MTB genome are differentially expressed and associated with six distinct transcriptional states as MTB enters into and exits from hypoxia. Moreover, there is strong evidence that the six-state model described in this paper relates to adaptations of MTB *in vivo*. The response to hypoxia is accompanied by other host-related stress mechanisms (e.g., alternative carbon utilization, iron limitation, copper stress), as a result of MTB’s evolutionary history as an intracellular pathogen. The large-scale expression changes demonstrate the importance of oxygen as a major force in the evolution of MTB and reveals that the pathogen alters gene expression in anticipation of future conditions and challenges. For example, the increased production of PE/PPE proteins during Late hypoxia in preparation for ESX export system production upon Resuscitation, thereby foreseeing the benefit of PE/PPE protein secretion for dissemination to other host cells. Integrating high-resolution and longitudinal profiling with experimentally determined TF-gene interactions enabled inference of key regulators and intricate circuit architecture that explain how the state transitions unfold (summarized in **Fig 4F**). Regulatory programs with characteristic motifs and properties were identified that serve to incorporate robustness (e.g., time-delay ensures state transition only upon proper conditions), synchronization (e.g., fold-change detection that uniforms response across all cells), and coordination across the states (e.g., interlocking FFLs, bifunctional Rv0081, mutual inhibition) as MTB transitions across time and O_2_ gradient. This study reveals that MTB encodes abundant network motifs, presumably with functions that cannot be carried out by simpler circuits, to successfully tailor MTB physiology to stresses within the host environment. It is interesting to speculate that these regulatory interactions have evolved in MTB as an adaptive response to ineffective immunity and failure to clear the pathogen. One of the most important challenges for antibiotic research will be to overcome these overlapping and redundant regulatory mechanisms with novel combinatorial interventions. This study presents significant steps toward apprehending these genetic programs in MTB, paving the way for predictive and rational strategies to improve clinical outcomes of TB treatment.

## Data and Software Availability

The datasets and computer code produced in this study are available in the following databases:

Hypoxia/reaeration RNA-Seq data: Gene Expression Omnibus GSE116353 https://www.ncbi.nlm.nih.gov/geo/query/acc.cgi?acc=GSE116353

The TF-target gene network for FanMod: MTB Network Portal Data Center http://networks.systemsbiology.net/mtb/data-center

R notebook with scripts for performing computation analyses: GitHub https://github.com/baliga-lab

## Acknowledgements

We thank members of the Baliga (in particular Christopher Plaisier now at Arizona State University) and Shmulevich labs for critical discussions, Lee Hiner (Airgas) for help with mass flow controllers, and the Center for Global Infectious Disease Research at Seattle Children’s Research Institute for maintenance and access to the BSL3 facilities. Funding was provided by the National Institute of Allergy and Infectious Diseases of the National Institutes of Health: pilot project grant awarded to Christopher Plaisier and E.J.R.P through U19AI135976; and R01AI128215; and U19AI10676. Funding was also provided by the National Science Foundation (NSF DBI-1565166).

## Author contributions

A.A., E.J.R.P., M.A-O., B.A., I.S. and N.S.B. designed research; A.A., E.J.R.P, M.P., A.K., and V.S. performed hypoxia experiments; A.A., E.J.R.P., M.A-O., J.Y. and B.A. analyzed data and performed computational analyses; A.A., E.J.R.P, M.A-O and N.S.B. wrote the paper.

## Competing interests

The authors declare no competing financial interests.

## Methods

The approaches used in this study include both computational and biological methods. Plots were generated using Python and R, and images prepared using Adobe Illustrator CS6 and Inkscape 0.91.

### Culturing conditions

Experiments were performed using H37Rv grown at 37°C in Middlebrook 7H9 supplemented with ADC and 0.05% Tween in spinner flasks. Working stocks were expanded from frozen aliquots shortly before experiments began. For hypoxia time-course experiment, a 50 mL culture was grown to mid-log phase, and diluted in 700 mL 7H9 media within each bioreactor to a starting A600 of 0.01. Cultures were stirred over a range of speeds throughout the experiment.

### Controlled O_2_ model design and operation

An Oxygen Sensor Spot (PreSens, Regensburg, Germany) was adhered within a 1L disposable spinner flask with two side arms (Corning, Corning, NY) using vacuum tweezers (Excelta, Buelton, CA). A velcro belt with a screw-on port for the fiber optic cable was wrapped around the flask. A gas line input was fastened on one arm of the flask, and a luer-lock/filter sampling port was connected to the other arm. Air and N_2_ gas lines were run into the Biological safety laboratory and connected to gas-specific mass flow controllers (Alicat Scientific, Tucson, AZ), whose outputs were connected downstream through a Y-connector that led into an incubator. Three separate flasks, all prepared as described above, were placed onto a stir plate inside an incubator at 37° C. The mixed gas line was split via additional Y-connecters, streamed through 0.2 um filters, and attached to the gas line inputs of each flask. Media was incubated overnight and checked for contamination before inoculated with MTB.

The mass flow controllers and oxygen sensor were linked to a computer, which could be remotely accessed and monitored in real-time. After inoculation, we programmed the mass flow controllers using Flow Vision software (Alicat Scientific) to achieve a changing gas mixture gradient, which allowed us creating a steady two-day depletion, followed by two-days of sustained hypoxia, and reaeration by flowing pure air into the headspace of the vessels and increasing the speed of the stir bars in each vessel.

### RNA isolation

Samples were collected by a luer-lock syringe to the sampling port. Sample volumes varied from 5 mL to 25 mL across the time course, depending on OD but were consistent across replicates of a time point. Samples were centrifuged at high speed for 5 min, supernatant was discarded and cell pellet was immediately flash frozen in liquid nitrogen. Cell pellets were stored at −80° C until all samples collected and then resuspended in 600 μL of fresh lysozyme solution in TE pH 8.0 (5 mg/mL). The resuspended cells were transferred to a tube containing Lysing Matrix B (MP Biomedicals, Santa Ana, CA) and incubated at 37° C for 30 min. Following incubation, 60 μL (1/10^th^ volume of lysate volume) of 10% SDS was added and then tubes were vigorously shaken at max speed for 30 s in a FastPrep 120 homogenizer (MP Biomedicals) three times. Tubes were centrifuged for 1 min (max speed), then 66 μL of 3 M sodium acetate pH 5.2 added and mixed well. Acid phenol (pH 4.2) was added at 726 μL and tubes were inverted to mix well (∼60 times). Samples were incubated at 65° C for 5 min, inverting tubes to mix samples every 30 s. Then, centrifuged at 14000 rpm for 5 min and upper aqueous phase was transferred to a new tube. 3M sodium acetate (pH 5.2) was added at 1/10^th^ volume along with 3x volumes of 100% ethanol. Sample was mixed well and incubated at −20° C for 1 hr or overnight. Following incubation, samples were centrifuged at 14000 rpm for 30 min at 4° C, ethanol was discarded and 500 μL of 70% ethanol was added. Samples were centrifuged again at 14000 rpm for 10 min at 4° C, supernatant discarded, and any residual ethanol removed using pipet. Pellet was allowed to air dry, resuspended in 30-40 μL of RNase free water and quantified by Nanodrop (Thermo Scientific). This was followed by in solution genomic DNA digestion using RQ1 Dnase (Promega) following manufacturer’s recommendation. RNA quality was analyzed in a 2100 Bioanalyzer system (Agilent Technologies). Total RNA samples were depleted of ribosomal RNA using the Ribo-Zero Bacteria rRNA Removal Kit (Illumina).

### Processing and analysis of RNA-seq data

Sample collection and RNA-extraction was performed as described above. Quality and purity of mRNA samples was determined with 2100 Bioanalyzer (Agilent, Santa Clara, CA). Samples were prepared with TrueSeq Stranded mRNA HT library preparation kit (Illumina, San Diego, CA) and multiplexed into a single run. All samples were sequenced on the NextSeq sequencing instrument in a high output 150 v2 flow cell. Paired-end 75 bp reads were checked for technical artifacts using Illumina default quality filtering steps. Raw FASTQ read data were processed using the R package DuffyNGS as described previously (Vignali et al., 2011). Briefly, raw reads were passed through a 3-stage alignment pipeline: (i) a prealignment stage to filter out unwanted transcripts, such as rRNA, mitochondrial RNA, albumin, and globin; (ii) a main genomic alignment stage against the genome(s) of interest; and (iii) a splice junction alignment stage against an index of standard and alternative exon splice junctions. Reads were aligned to *M. tuberculosis H37Rv* (ASM19595v2) with Bowtie2 (Langmead and Salzberg, 2012), using the command line option “very-sensitive.” BAM files from stages (ii) and (iii) were combined into read depth wiggle tracks that recorded both uniquely mapped and multiply mapped reads to each of the forward and reverse strands of the genome(s) at single-nucleotide resolution. Gene transcript abundance was then measured by summing total reads landing inside annotated gene boundaries, expressed as both RPKM and raw read counts. Two stringencies of gene abundance were provided using all aligned reads and by just counting uniquely aligned reads.

### Differential expression

We used the raw read counts, estimated with DuffyNGS as described above, as input for DESeq2 (Love et al., 2014). We compared the transcriptional profile of each time point respect to T0. Genes with adjusted *P*-value < 0.05 and estimated absolute log2 fold-change >1 were considered differentially expressed.

### Identification of transcriptional states adopted by MTB in the controlled O_2_ model

After normalizing the full transcriptional dataset using DESeq2, we used Principal Component Analysis (PCA) to reduce the dimensionality of our data (median transcript levels of 4,049 genes profiled at each time point). Then, we used the NbClust R package (Charrad et al., 2014) to identify the most likely number of clusters in the PCA space. Two clusters were predicted (**Fig S2A**). One cluster contained the time points: T8, T9, T11, T19, T20 and T21. A second cluster contained the other time points. Motivated by the clustering of consecutive time points and their similarity in O_2_ concentrations in **Fig S2A** (T0-T1, T2-T7,T16-T18, etc), we evaluated the structure of the largest cluster using PCA and NbClust as before. However, this time the T8, T9, T11, T19, T20 and T21 points were removed before the analysis. Five distinct clusters were identified (**Fig S2B**). To confirm the presence of six clusters (each corresponding to a transcriptional state), we performed hierarchical clustering of the transcriptional dataset with bootstrapping using the Pvclust R package (Suzuki and Shimodaira, 2006) (**Fig S2C**). All six putative clusters had 100% bootstrap support in the resulting dendrogram. Finally, we confirmed the presence of the six defined transcriptional states with the t-distributed stochastic neighbor embedding (tSNE) algorithm (**Fig 2A**).

### Connecting differentially expressed genes with the six hypoxia-related transcriptional states of MTB

To understand the functional implications of the transcriptional states adopted by MTB during entry and exit from hypoxia, each differentially expressed gene was assigned to the state in which it had the highest average transcription level. As an unsupervised alternative, we used the Boruta R package (Kursa, 2010), that implements random forest to select all features (in our case transcriptional profiles), to identify the genes that distinguish any given state from the rest. There was statistically significant overlap between the groups of genes associated to any given transcriptional state by the two approaches (**Table S4**). Because Boruta only selected 566 genes (out of 2,582 differentially expressed genes), we decided to use the average transcriptional profile based gene assignment. In this way we tried to capture the biological processes active in the different states without excluding any differentially expressed gene. To evaluate the quality of the resulting sets of genes, we computed the mean square residual (MSR) of each gene cluster (**Table S4**). The MSR is widely used as a metric of performance of biclustering methods (which cluster both genes and conditions) (Reiss et al., 2006). A low MSR value indicates that individual profiles do not deviate from the average profile of the bicluster (in our case, the group of genes in the relevant time points/state). We also computed the mean Pearson correlation among the genes assigned to each transcriptional state (**Table S4**). In support of our gene assignment, the sets of genes associated to MTB transcriptional states had MSR values smaller than 0.1 In fact, for the Depletion and Late hypoxia genes the MSR values were smaller than 0.05. The average Pearson correlation values were equal to or greater than 0.4 for all gene clusters but Normoxia.

### Metabolic pathway analysis

We mapped the measured gene expression data against the most recent genome-scale metabolic network construction of *M. tuberculosis* H37Rv iEK1011 (Kavvas et al., 2018) using COBRApy (Ebrahim et al., 2013). We used the subsystem defintions defined in iEK1011 to explore pathway usage at the network level.

### MTB ChIP-seq derived TF-gene network

The initial ChIP-seq derived MTB network consisted of 6,581 interactions occurring in the −150bp to +70bp region of genes’ promoter reported by Minch et al. (Minch et al., 2015). We expanded that MTB ChIP-seq network by taking into account operon organizations. For a given TF-gene interaction, if the target gene is part of an operon, we included all other members of the operon as potential targets of the corresponding TF. The expanded MTB ChIP-seq network contained 12,188 interactions.

### Detection of network motifs in the MTB ChIP-seq network

To identify network motifs in the MTB transcriptional network, we used the MotifNet webserver (Smoly et al., 2017). We scanned for all potential three and four nodes motifs with maximum *P*-value ≤ 0.01 (in 1000 random networks) and with 100 or more instances in the analyzed network. For this analysis, we constrained the ChIP-seq derived network by excluding genes that were not differentially expressed in our time course. This network filtering was done to improve detection of motifs most relevant to the actual changes in transcript levels we observed.

### DREM analysis

DREM2.0 (Schulz et al., 2012) was run with default parameters. The input TF-gene network was the MTB ChIP-seq network described above. The input expression data contained the median transcriptional profiles of the 2,582 differentially expressed genes. The minimum absolute expression change parameter was set to 0.75.

### Permutation test for evaluating significance of overlap between TF regulons and sets of genes associated with identified transcriptional states

The 2,582 differentially expressed genes in the controlled O_2_ model were permutated 1000 times to generate shuffled gene clusters (corresponding to the six transcriptional states). In each permutation, the produced shuffled gene clusters had the same size as the original ones. Then, significance of the overlap between regulons of differentially expressed TFs and the shuffled gene sets was evaluated using a hypergeometric test. Hypergeometric test *P*-values below 0.05 were considered significant. The overall permutation test *P*-value was computed as the proportion of cases (out of 1000 permutations) in which the number of enriched regulons was equal or higher than the observed values in the original data.

### Differential expression analysis of transcriptional data collected with the defined hypoxia model

We downloaded the transcriptional profile data of MTB at day 0, day 1, day 3, day 5, day 7 and day 8 (reaeration) collected by Galagan et al (Galagan et al., 2013) using the defined hypoxia model (GEO accession number: GSE43466). We performed a Bayesian t-test using CyberT (Baldi and Long, 2001) to compare the gene expression profiles at each time point respect to T0. Genes with adjusted *P*-value < 0.05 and absolute log2 fold-change > 1 were considered differentially expressed.

### Modeling of Rv0081-directed I-FFL with Late hypoxia gene expression

The IFFL motif is commonly modeled by the following equations (Goentoro et al., 2009):

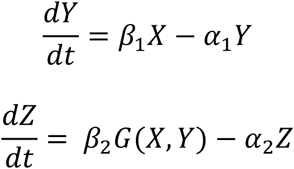

where *Z* is the output of the motif which in our case represents the average expression of Late hypoxia genes between T7-T20. *X* and *Y* represent the expression of Rv0081 and average expression of Early hypoxia TFs (excluding Rv0081), respectively. Goentoro et al (Goentoro et al., 2009) analyzed three possible models of the I-FFL motif representing three configurations in which Late hypoxia genes could be regulated. Their focus was to identify conditions for which the I-FFL motif works as fold change detection. The models correspond to exclusive binding, independent binding, and cooperative binding which are represented by the following equations (Goentoro et al., 2009):

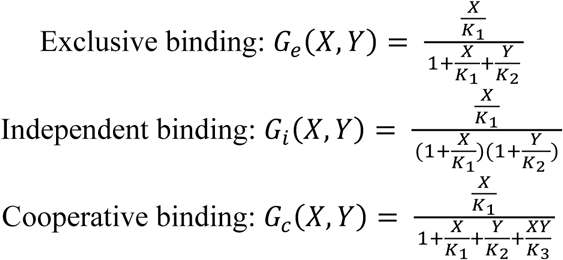

where *G*_*e*_, *G*_*i*_, and *G*_*c*_ are functions that determine the rate change of Late hypoxia genes (*Z*); *K*_*1*_ is the binding rate between Rv0081 and Late hypoxia genes; *K*_*2*_ is the binding rate between Early hypoxia TFs and Late hypoxia genes; and *K*_*3*_ is the cooperative binding rate of Rv0081 and Early hypoxia TFs with the Late hypoxia genes. *α*_*2*_, *β*_*2*_, *K*_*1*_, *K*_*2*_, and *K*_*3*_ were estimated by an optimization procedure using the average experimental values of *X, Y, Z* at different time points. We used the Nelder-Mead simplex algorithm for optimization (Nelder, 1965) as implemented in MATLAB R2014a. The objective function used for minimization is the root-mean-squared deviation (RMSD) between experimental and estimated values of Late hypoxia gene expression is given by:

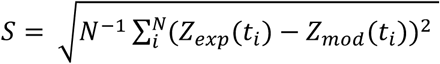

where *Z*_*exp*_(*t*_*i*_) is the average expression of Late hypoxia genes at time *t*_*i*_ obtained experimentally, and *Z*_*mod*_(*t*_*i*_) is the corresponding model estimate. Similarly, *α*_*1*_ and *β*_*1*_ were estimated by a similar optimization procedure using the average experimental values of *Y* and *X*. The objective function in this case is given by:

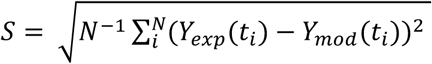

where *Y*_*exp*_(*t*_*i*_) is the average expression of repressors at time *t*_*i*_ obtained experimentally, and *Y*_*mod*_(*t*_*i*_) is the corresponding model estimate.

## Supplemental Materials

**Figure S1.**
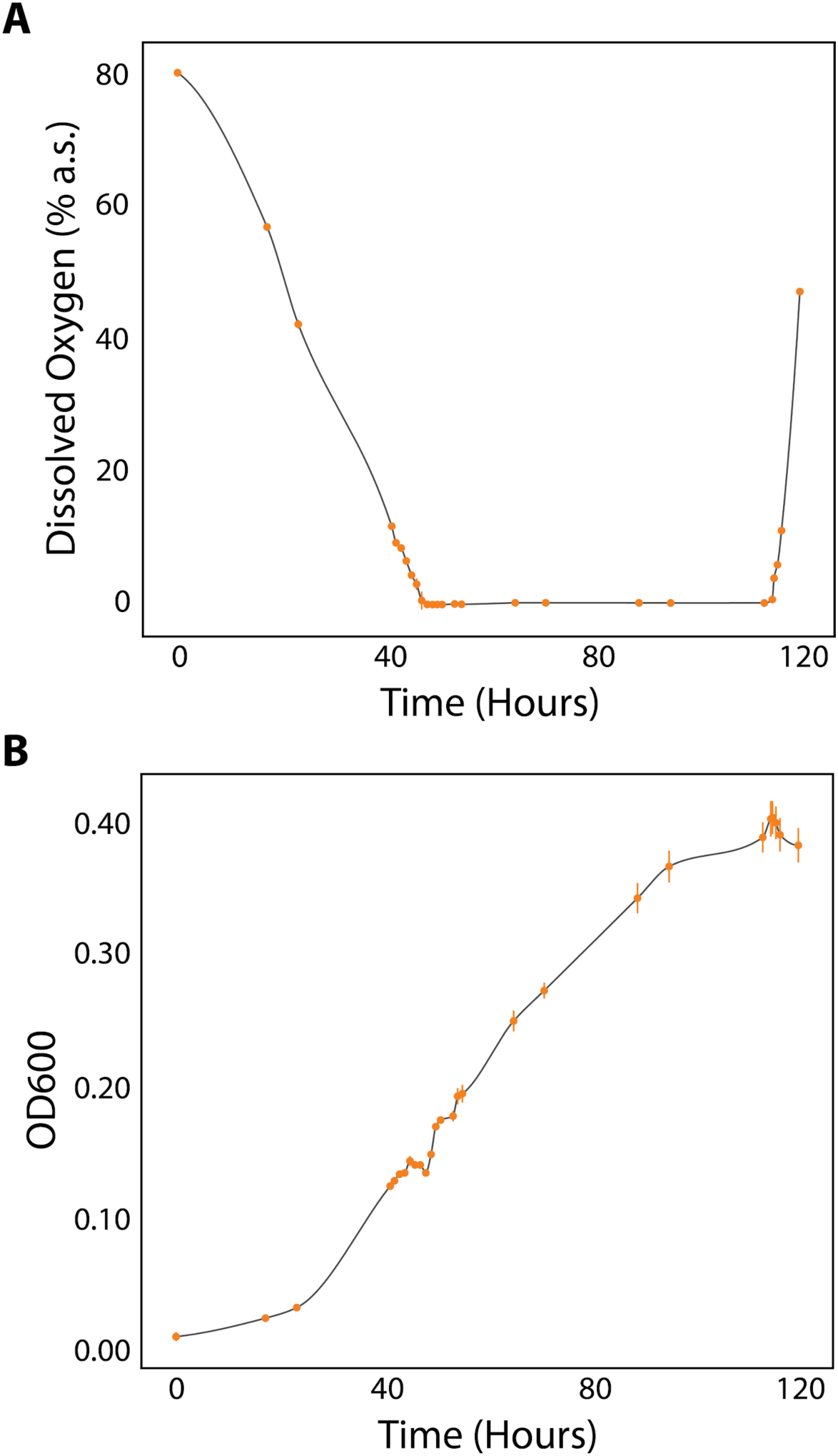
Bioreactor system is highly reproducible across three biological replicates. **(A)** Dissolved oxygen curve of growth medium across hypoxia and reaeration time-course measured with oxygen sensor spots (Presens). **(B)** Optical density across replicates over the time course. Datapoints are the median of three replicates with error bars showing the Standard Error of measurements.

**Figure S2.**
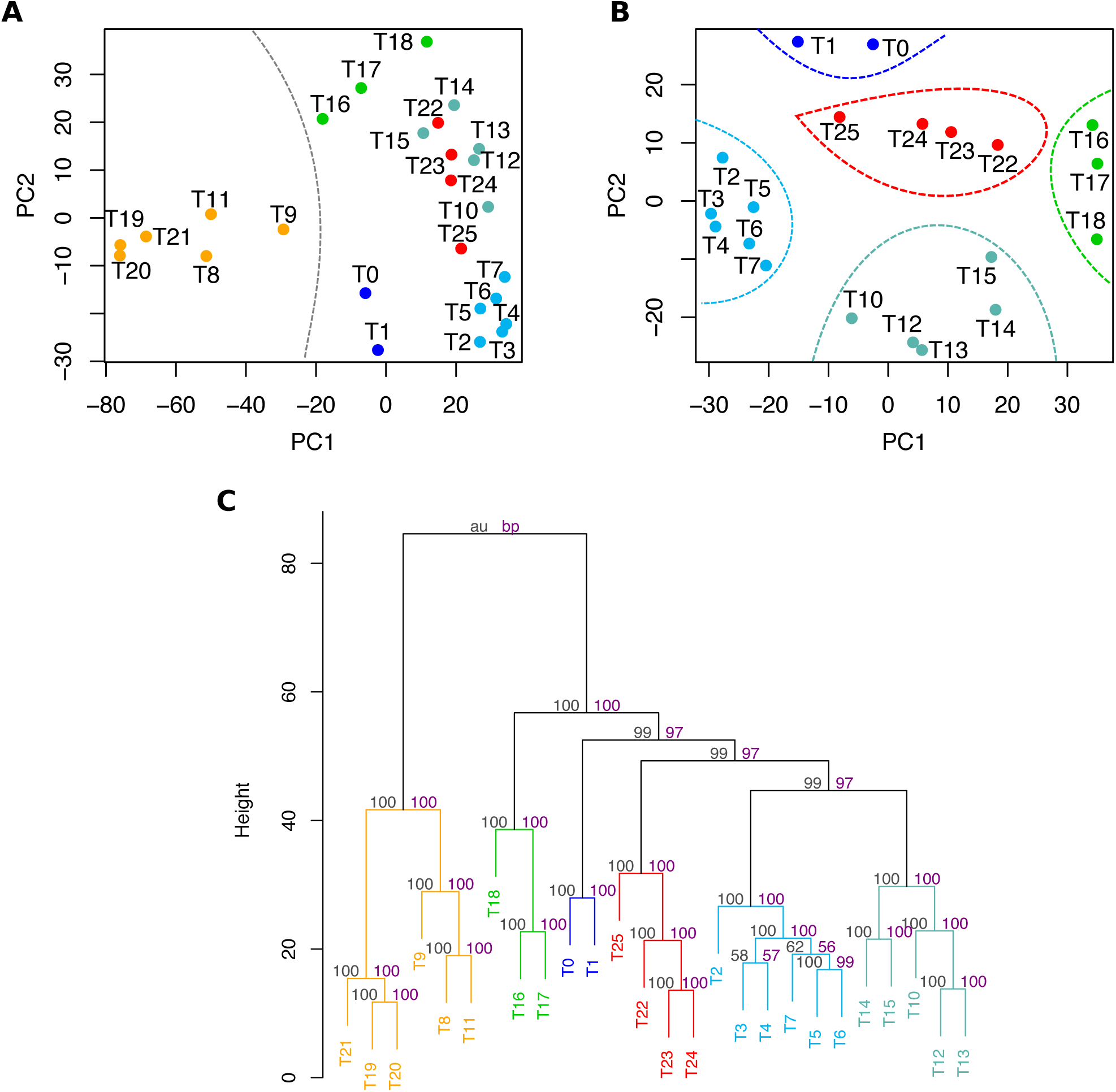
Identification of six transcriptional states of MTB during depletion and re-introduction of O_2_ in the controlled O_2_ model. Time points and dendrogram branches are color coded according to their membership in the six transcriptional states defined in **Fig 2A**. (**A**) Principal component plot of the transcriptional matrix containing the median profiles of all 4,049 genes. Dashed line indicates the two clusters identified by the NbClust tool in R (Charrad et al., 2014). (**B**) Principal component plot of the same transcriptional matrix after excluding T8, T9, T11, T19-T21 data. Dashed lines indicate the five clusters identified by the NbClust tool in R. (**C**) Hierarchical clustering of the transcriptional data using the Pvclust R package (Suzuki and Shimodaira, 2006). Numbers on top of the dendrogram branches indicate the unbiased *P*-value (au; black) and bootstrap support (bp; purple) for each branch.

**Figure S3.**
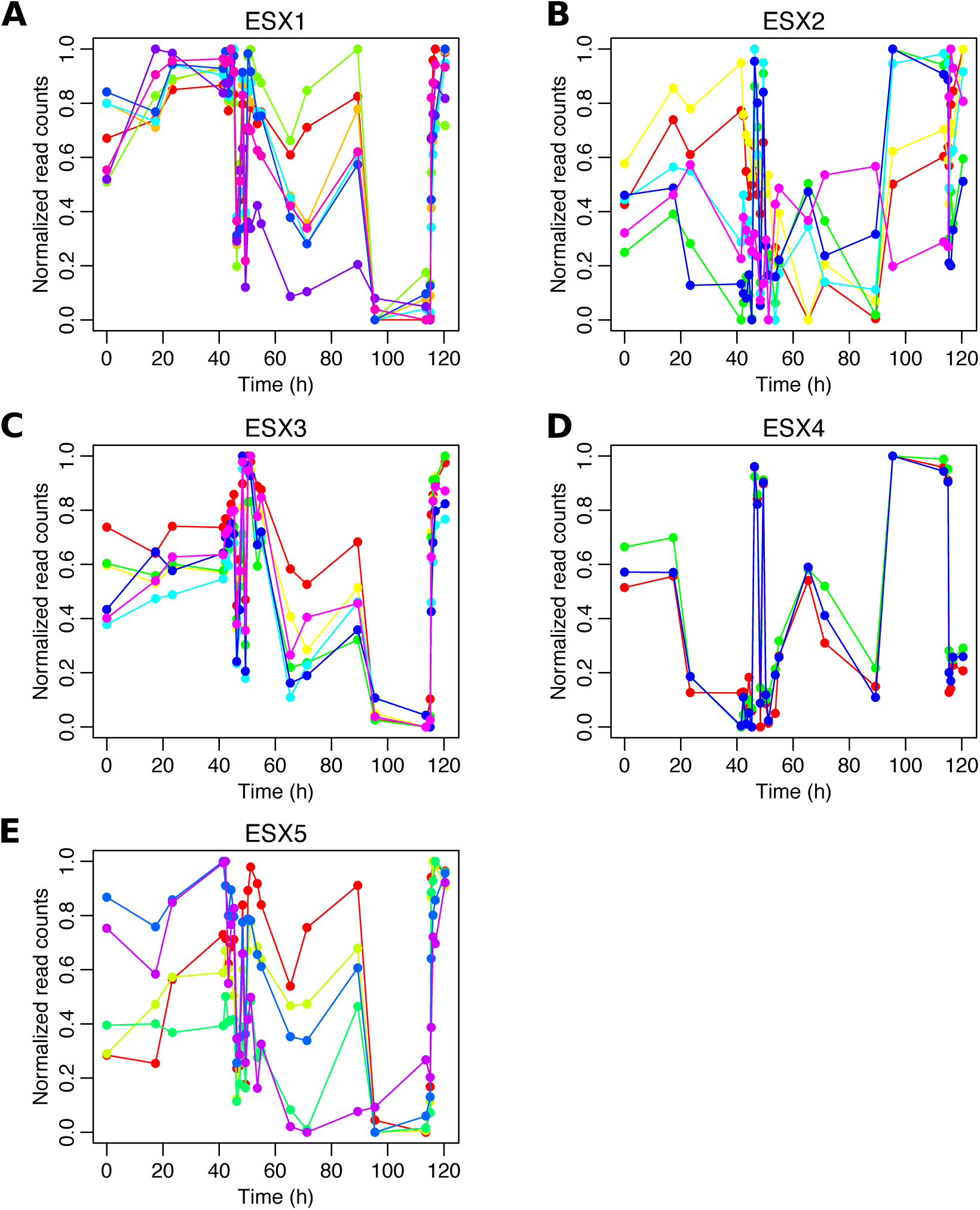
Transcriptional profile of the five type seven secretion systems (T7SS) in MTB. The transcriptional profiles of genes encoding components of MTB’s ESX systems (also known as T7SS) are shown. Each gene is represented with a color. For visualization purpose, the normalized read counts were re-scaled to the [0,1] interval.

**Figure S4.**
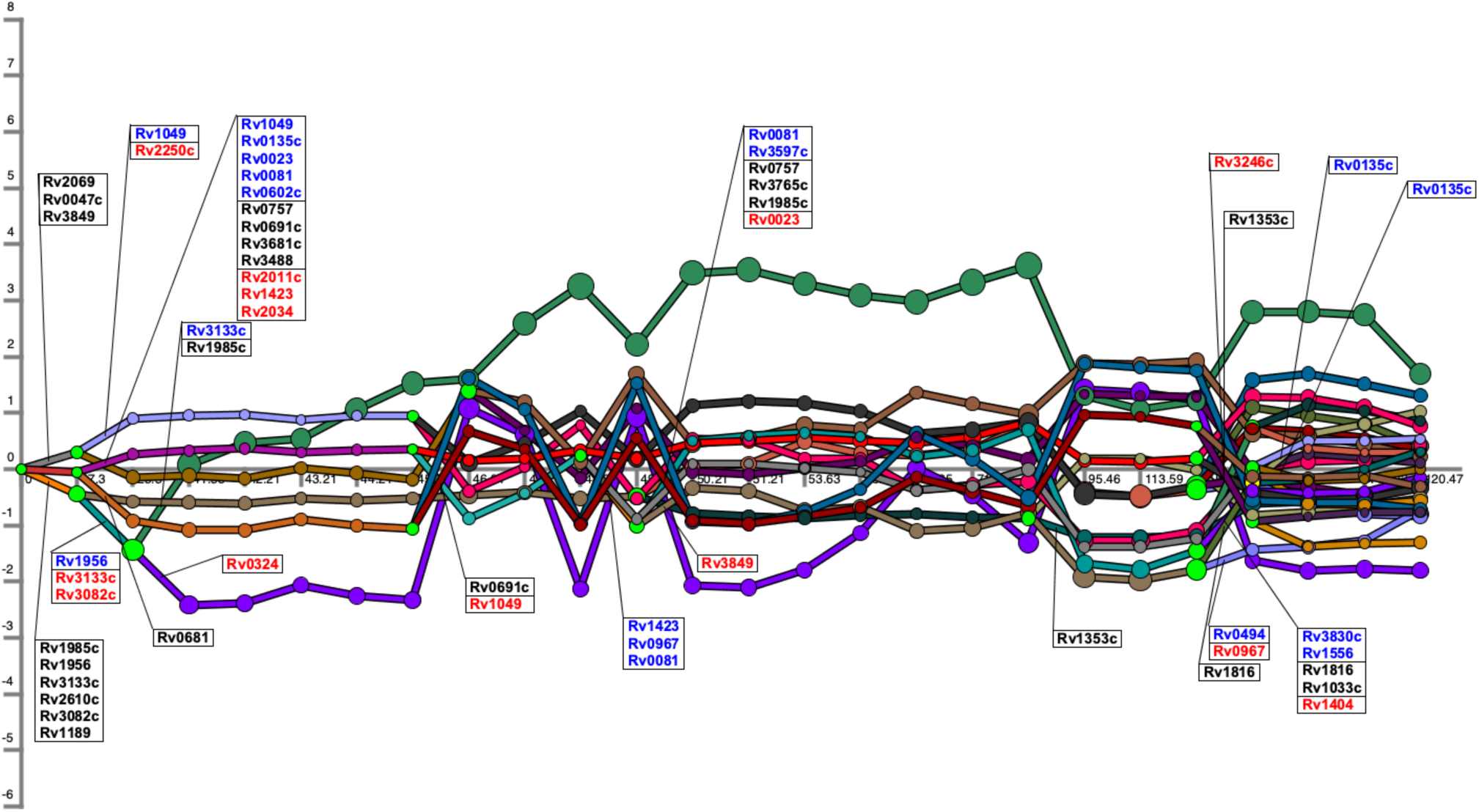
DREM output. DREM associated 35 TFs with the 23 identified branching points (light green nodes). Blue, black and red TFs labels indicate up-regulation, no change or down-regulation, respectively.

**Figure S5.**
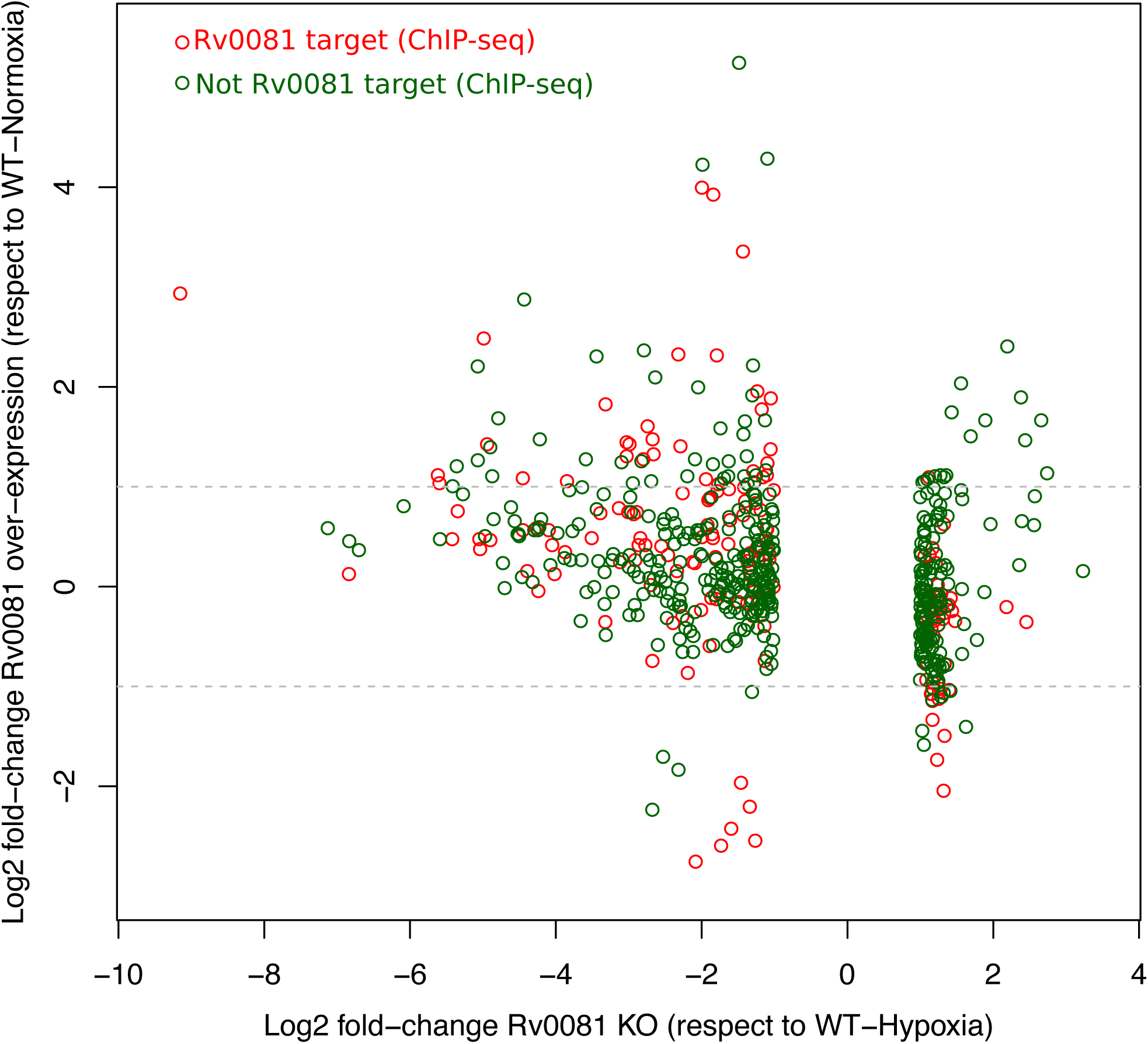
The dual activity of Rv0081. Plot is based on the comparison between the normoxic TFOE data and hypoxic ΔRv0081 data (Rustad et al., 2014; Sun et al., 2018). Only genes with differential expression in the Rv0081deletion strain during hypoxia are shown. This set of genes was separated in the ones that are controlled by Rv0081 (based on ChIP-seq data) and the ones that are not.

**Figure S6.**
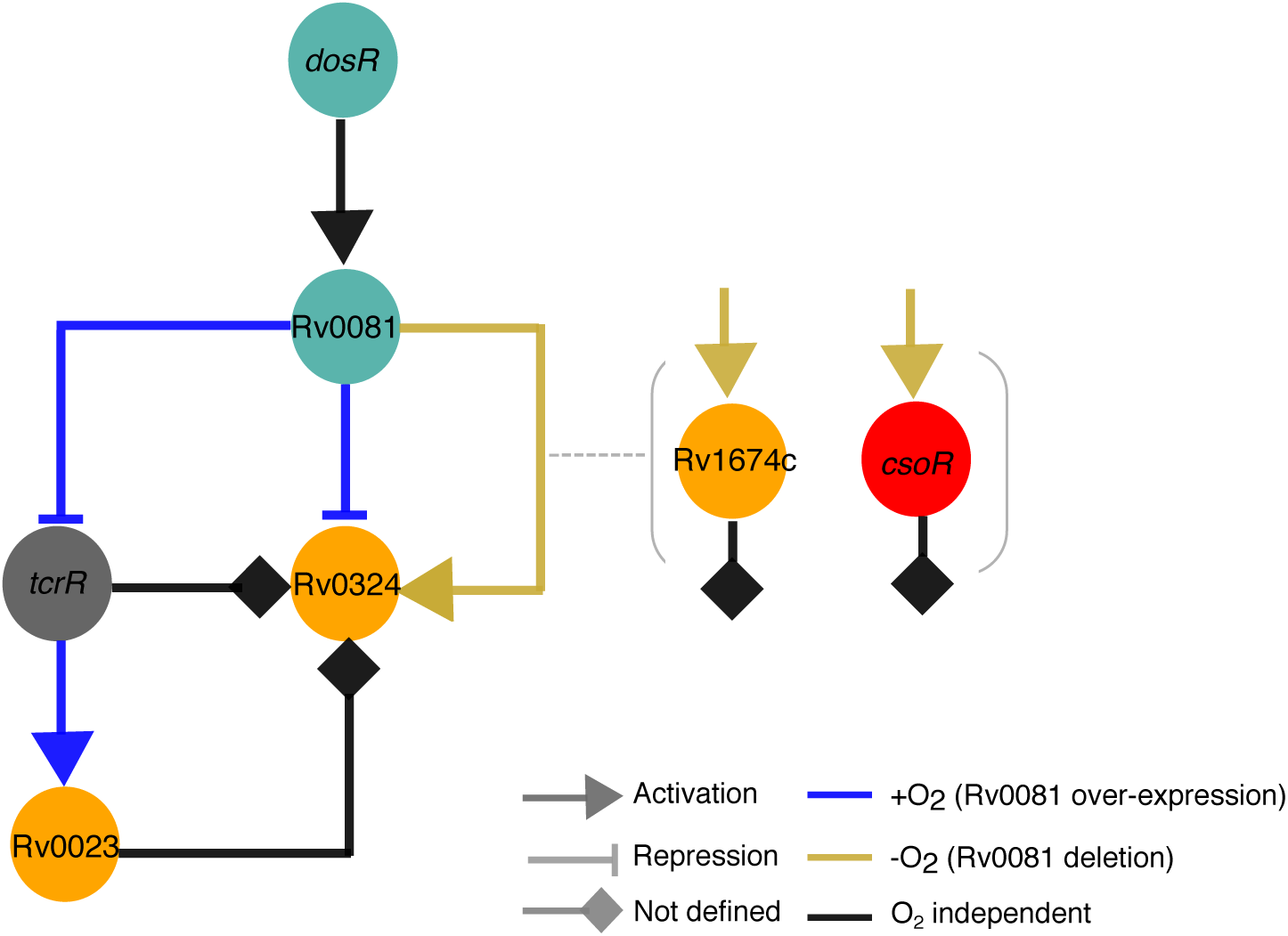
Dual activity of Rv0081 seems to be necessary for coordinating the repression and activation of Rv0324 at early and late points of the time course. Nodes are colored based on their transcriptional states membership. The shown transcriptional circuit was derived from available protein-DNA biding data in MTB (Minch et al., 2015). The signs of the TF-TF interactions were defined using the change in transcript levels of the target genes in the TFOE strains (blue edges) and hypoxic ΔRv0081 strain (golden edges) (Rustad et al., 2014; Sun et al., 2018). Interactions with no assigned signs are shown in black.

**Figure S7.**
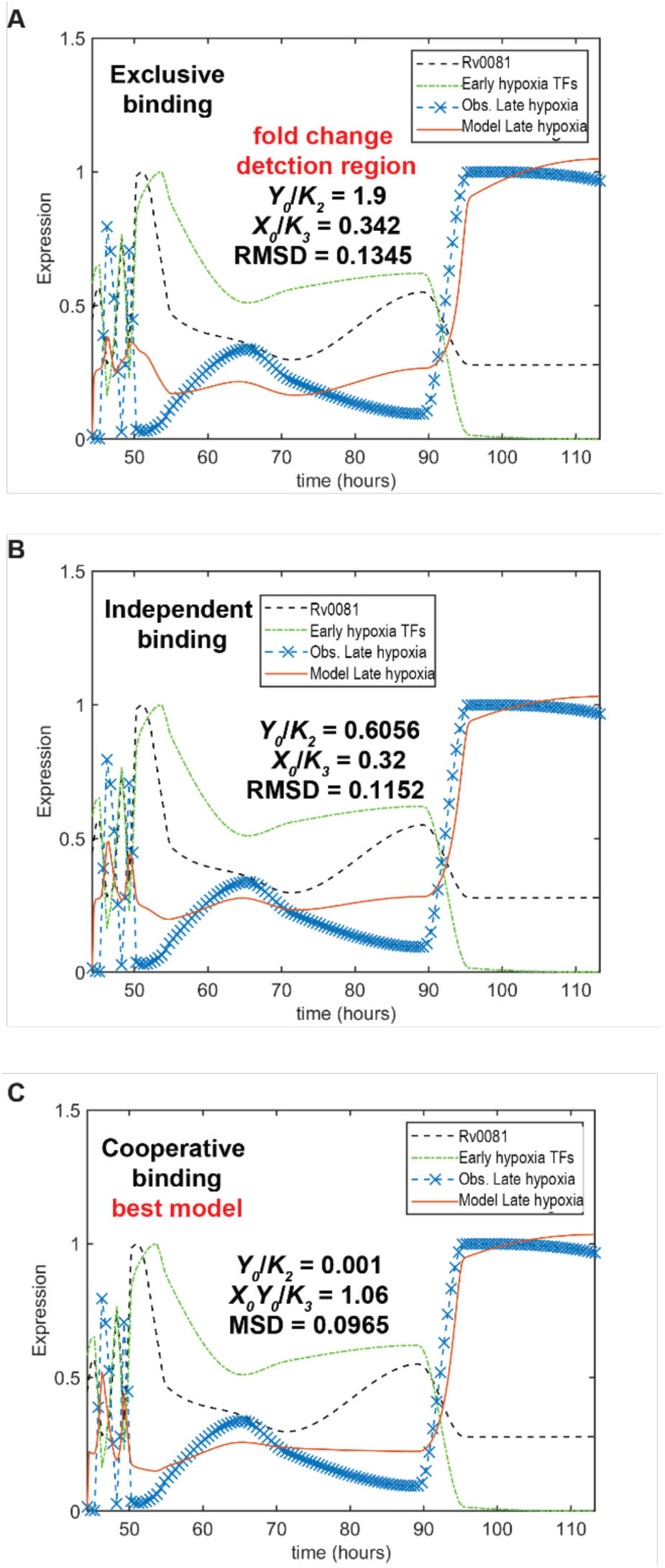
Modeling of an I-FFL controlling the expression of Late hypoxia genes using methods of Goentoro et al (Goentoro et al., 2009). Simulation results for the I-FFL motif under (A) exclusive, (B) independent, and (C) cooperative binding configurations (*see methods*). *X*_*o*_ and *Y*_*o*_ represent the observed expression of Rv0081 and average observed expression of Early hypoxia TFs (excluding Rv0081), respectively. *K*_*1*_ is the binding rate between Rv0081 and Late hypoxia genes; *K*_*2*_ is the binding rate between Early hypoxia TFs and Late hypoxia genes; and *K*_*3*_ is the cooperative binding rate of Rv0081 and Early hypoxia TFs with the Late hypoxia genes. RMSD is the root-mean-squared deviation between experimental and estimated values of Late hypoxia gene expression.

**Table S1.** Gene set overlap between the three models of hypoxia-induced dormancy.

| State | IHR<br>recall* | Overlap<br>*P*-value | EHR<br>recall<sup>\$</sup> | Overlap<br>*P*-value | NRP1<br>recall<sup>%</sup> | Overlap<br>*P*-value | NRP2<br>recall<sup>^</sup> | Overlap<br>*P*-value |
| --- | --- | --- | --- | --- | --- | --- | --- | --- |
| Normoxia | 0 | 1 | 0.004 | 0.992 | 0.016 | 0.739 | 0.004 | 0.991 |
| Depletion | 0.02 | 0.997 | 0.035 | 1 | 0.107 | 0.597 | 0.04 | 1 |
| Early hypoxia | 0.408 | 2.9E-10 | 0.070 | 0.783 | 0.176 | 0.116 | 0.138 | 1.9E-03 |
| Mid hypoxia | 0.49 | 1.7E-14 | 0.226 | 6.3E-13 | 0.187 | 8.3E-07 | 0.201 | 1.8E-09 |
| Late hypoxia | 0.041 | 1 | 0.196 | 0.963 | 0.032 | 1 | 0.125 | 1 |
| Resuscitation | 0.02 | 0.996 | 0.165 | 3.2E-03 | 0.316 | 8.6E-16 | 0.214 | 8.2E-07 |

\* Initial Hypoxic Response (IHR) contains 49 genes (Rustad et al., 2008)
<sup>\$</sup> Enduring Hypoxic Response (EHR) contains 230 genes (Rustad et al., 2008)
<sup>%</sup> Non Replicating Persistence 1 (NRP1) contains 187 genes (Muttucumaru et al., 2004)
<sup>^</sup> Non Replicating Persistence 2 (NRP2) contains 224 genes (Muttucumaru et al., 2004)

**Table S2.**
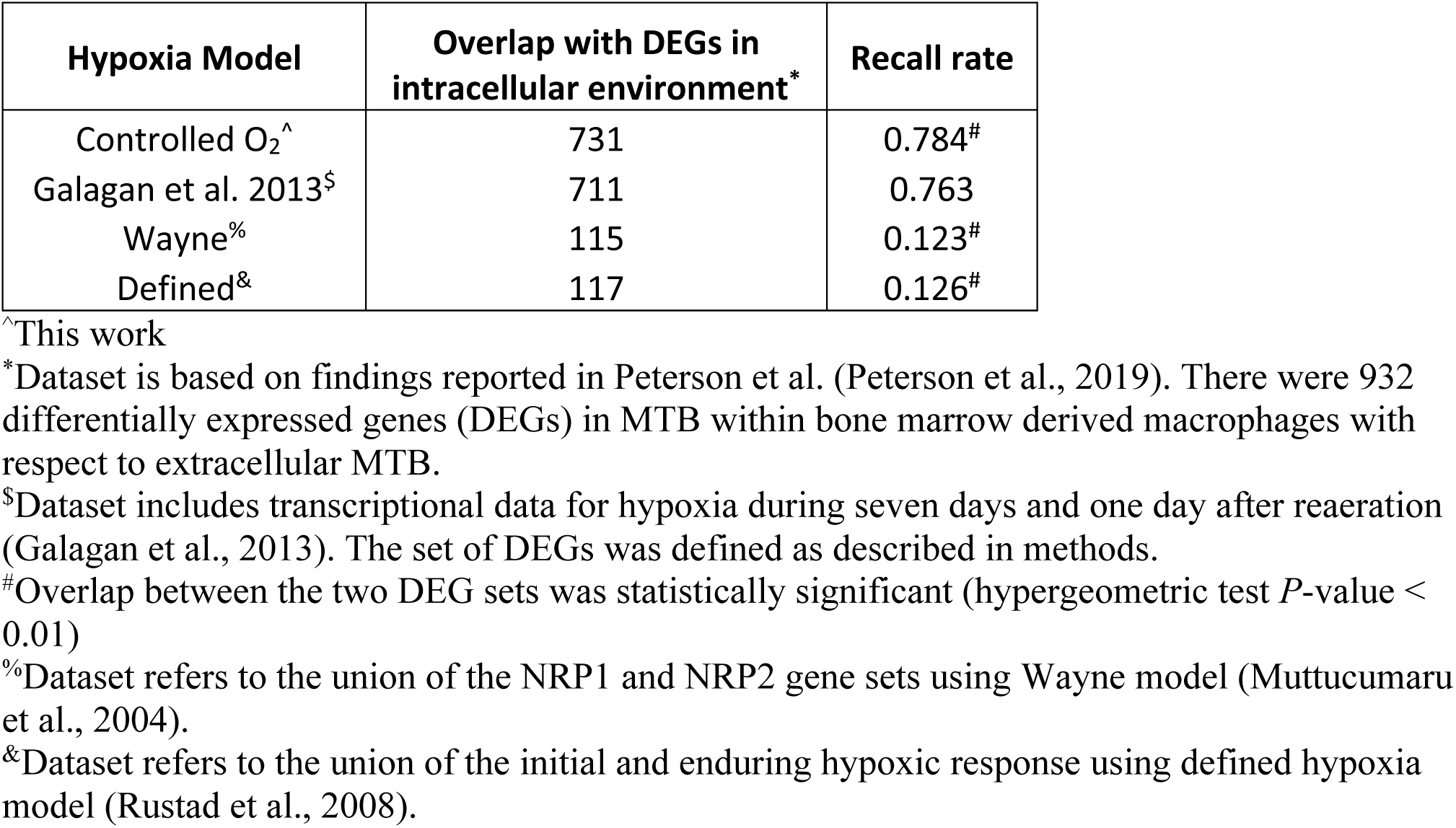
Gene set overlap between the models of hypoxia-induced dormancy and intracellular MTB dataset.

| Hypoxia Model | Overlap with DEGs in intracellular environment* | Recall rate |
| --- | --- | --- |
| Controlled O<sub>2</sub><sup>^</sup> | 731 | 0.784<sup>#</sup> |
| Galagan et al. 2013<sup>\$</sup> | 711 | 0.763 |
| Wayne<sup>%</sup> | 115 | 0.123<sup>#</sup> |
| Defined<sup>&</sup> | 117 | 0.126<sup>#</sup> |

<sup>^</sup>This work
<sup>\*</sup>Dataset is based on findings reported in Peterson et al. (Peterson et al., 2019). There were 932 differentially expressed genes (DEGs) in MTB within bone marrow derived macrophages with respect to extracellular MTB.
<sup>\$</sup>Dataset includes transcriptional data for hypoxia during seven days and one day after reaeration (Galagan et al., 2013). The set of DEGs was defined as described in methods.
<sup>#</sup>Overlap between the two DEG sets was statistically significant (hypergeometric test $P$ -value < 0.01)
<sup>%</sup>Dataset refers to the union of the NRP1 and NRP2 gene sets using Wayne model (Muttucumaru et al., 2004).
<sup>&</sup>Dataset refers to the union of the initial and enduring hypoxic response using defined hypoxia model (Rustad et al., 2008).

**Table S3.**
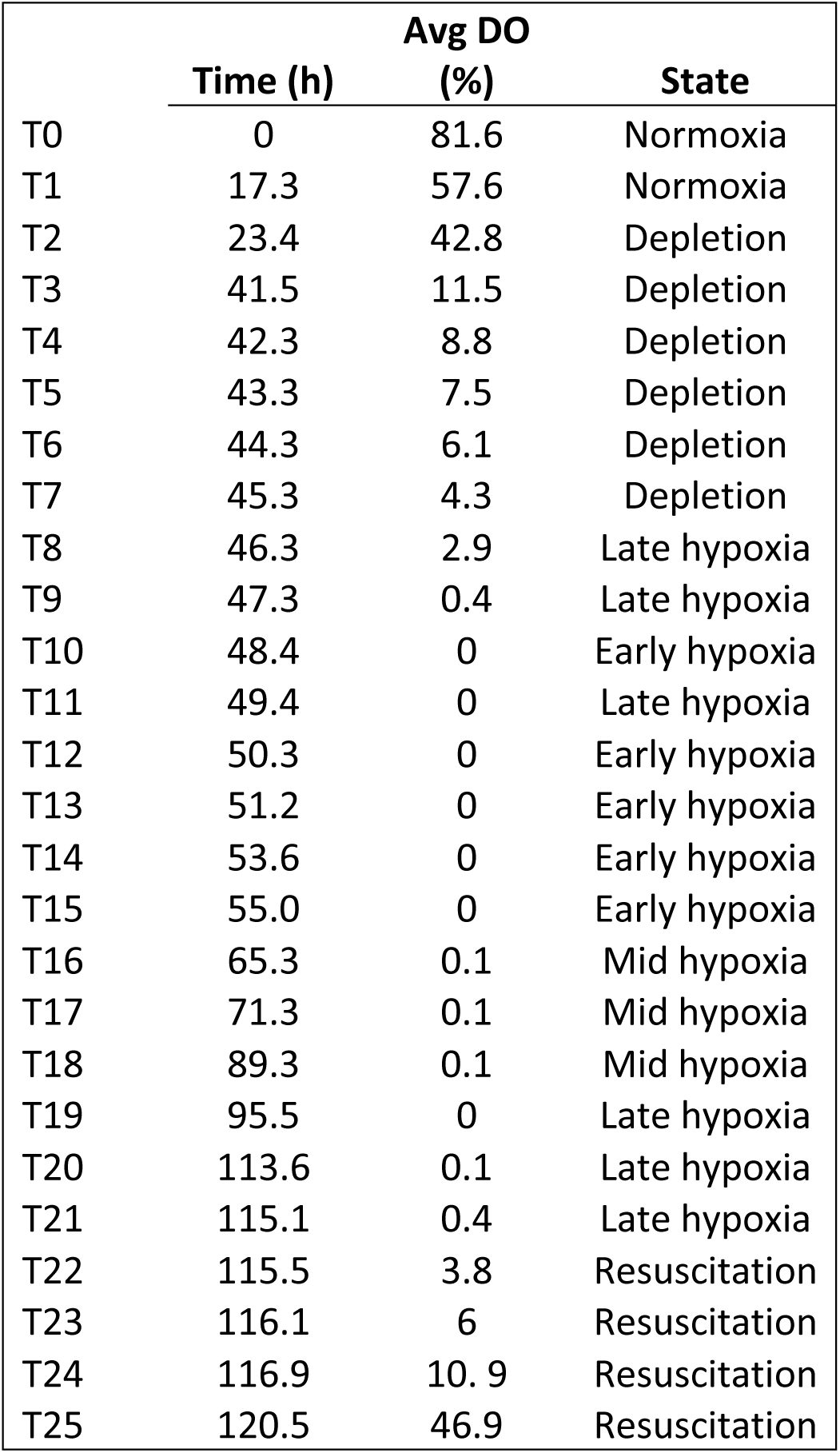
Samples (T0 to T25) were used to define the six state model. Each state was also defined with oxygen and time intervals.

|  | **Time (h)** | **Avg DO (%)** | **State** |
| --- | --- | --- | --- |
| T0 | 0 | 81.6 | Normoxia |
| T1 | 17.3 | 57.6 | Normoxia |
| T2 | 23.4 | 42.8 | Depletion |
| T3 | 41.5 | 11.5 | Depletion |
| T4 | 42.3 | 8.8 | Depletion |
| T5 | 43.3 | 7.5 | Depletion |
| T6 | 44.3 | 6.1 | Depletion |
| T7 | 45.3 | 4.3 | Depletion |
| T8 | 46.3 | 2.9 | Late hypoxia |
| T9 | 47.3 | 0.4 | Late hypoxia |
| T10 | 48.4 | 0 | Early hypoxia |
| T11 | 49.4 | 0 | Late hypoxia |
| T12 | 50.3 | 0 | Early hypoxia |
| T13 | 51.2 | 0 | Early hypoxia |
| T14 | 53.6 | 0 | Early hypoxia |
| T15 | 55.0 | 0 | Early hypoxia |
| T16 | 65.3 | 0.1 | Mid hypoxia |
| T17 | 71.3 | 0.1 | Mid hypoxia |
| T18 | 89.3 | 0.1 | Mid hypoxia |
| T19 | 95.5 | 0 | Late hypoxia |
| T20 | 113.6 | 0.1 | Late hypoxia |
| T21 | 115.1 | 0.4 | Late hypoxia |
| T22 | 115.5 | 3.8 | Resuscitation |
| T23 | 116.1 | 6 | Resuscitation |
| T24 | 116.9 | 10.9 | Resuscitation |
| T25 | 120.5 | 46.9 | Resuscitation |

**Table S4.** Properties of gene clusters associated with identified transcriptional states.

| State | Genes selected<br>based on<br>average | Mean<br>Square<br>Residual | Pearson<br>correlation | Boruta<br>genes | Overlap | Recall<br>Boruta<br>genes | *P*-value<br>overlap |
| --- | --- | --- | --- | --- | --- | --- | --- |
| Normoxia | 81 | 0.04 | 0.28 | 15 | 3 | 0.20 | 9.40E-05 |
| Depletion | 446 | 0.02 | 0.71 | 127 | 73 | 0.57 | 7.70E-27 |
| Early Hypoxia | 328 | 0.05 | 0.55 | 80 | 61 | 0.76 | 6.00E-42 |
| Mid Hypoxia | 320 | 0.08 | 0.4 | 44 | 24 | 0.55 | 1.40E-12 |
| Late Hypoxia | 978 | 0.03 | 0.75 | 254 | 185 | 0.73 | 1.70E-33 |
| Resuscitation | 429 | 0.09 | 0.47 | 94 | 56 | 0.60 | 8.10E-23 |

**Dataset S1**. Each cluster represents a distinct transcriptional state and was classified with sets of non-overlapping differentially expressed genes (see methods): Normoxia (81 genes), Transition (446 genes), Stage Ia hypoxia (328 genes), Stage Ib hypoxia (320 genes), Stage II hypoxia (978 genes), and Resuscitation (429 genes) (**Dataset S1**).

**Dataset S2**: TF regulons enrichment with genes associated with identified transcriptional states.

## REFERENCES

Abdallah, A.M., Savage, N.D., van Zon, M., Wilson, L., Vandenbroucke-Grauls, C.M., van der Wel, N.N., Ottenhoff, T.H., and Bitter, W. (2008). The ESX-5 secretion system of Mycobacterium marinum modulates the macrophage response. J Immunol 181, 7166–7175.

Alon, U. (2007). Network motifs: theory and experimental approaches. Nat Rev Genet 8, 450–461.

Baldi, P., and Long, A.D. (2001). A Bayesian framework for the analysis of microarray expression data: regularized t-test and statistical inferences of gene changes. Bioinformatics 17, 509–519.

Bartek, I.L., Woolhiser, L.K., Baughn, A.D., Basaraba, R.J., Jacobs, W.R., Jr., Lenaerts, A.J., and Voskuil, M.I. (2014). Mycobacterium tuberculosis Lsr2 is a global transcriptional regulator required for adaptation to changing oxygen levels and virulence. MBio 5, e01106–01114.

Baugh, L.R., Hill, A.A., Claggett, J.M., Hill-Harfe, K., Wen, J.C., Slonim, D.K., Brown, E.L., and Hunter, C.P. (2005). The homeodomain protein PAL-1 specifies a lineage-specific regulatory network in the C. elegans embryo. Development 132, 1843–1854.

Bolouri, H.Y. M.; Beilke, J.; Johnson, R.; Fox, B.; Huang, L.; Costa Santini, C.; Hill, C.M.; van der Vuurst de Vries, A-R.; Shannon, P.; Dervan, A.; Sivakumar, P.; Trotter, M.; Bassett, D.; Ratushny, A. (2019). Integrative network modeling reveals mechanisms underlying T cell exhaustion.

Boon, C., and Dick, T. (2002). Mycobacterium bovis BCG response regulator essential for hypoxic dormancy. J Bacteriol 184, 6760–6767.

Bottai, D., and Brosch, R. (2009). Mycobacterial PE, PPE and ESX clusters: novel insights into the secretion of these most unusual protein families. Mol Microbiol 73, 325–328.

Bromberg, K.D., Ma’ayan, A., Neves, S.R., and Iyengar, R. (2008). Design logic of a cannabinoid receptor signaling network that triggers neurite outgrowth. Science 320, 903–909.

Chao, M.C., and Rubin, E.J. (2010). Letting sleeping dos lie: does dormancy play a role in tuberculosis? Annu Rev Microbiol 64, 293–311.

Charrad, M., Ghazzali, N., Boiteau, V., and Niknafs, A. (2014). NbClust: An R Package for Determining the Relevant Number of Clusters in a Data Set. Journal of Statistical Software 61.

Cole, S.T., Brosch, R., Parkhill, J., Garnier, T., Churcher, C., Harris, D., Gordon, S.V., Eiglmeier, K., Gas, S., Barry, C.E., 3rd, et al. (1998). Deciphering the biology of Mycobacterium tuberculosis from the complete genome sequence. Nature 393, 537–544.

Consortium, E., Roy, S., Ernst, J., Kharchenko, P.V., Kheradpour, P., Negre, N., Eaton, M.L., Landolin, J.M., Bristow, C.A., Ma, L., et al. (2010). Identification of functional elements and regulatory circuits by Drosophila modENCODE. Science 330, 1787–1797.

Cortes, T., Schubert, O.T., Banaei-Esfahani, A., Collins, B.C., Aebersold, R., and Young, D.B. (2017). Delayed effects of transcriptional responses in Mycobacterium tuberculosis exposed to nitric oxide suggest other mechanisms involved in survival. Sci Rep 7, 8208.

Ebrahim, A., Lerman, J.A., Palsson, B.O., and Hyduke, D.R. (2013). COBRApy: COnstraints-Based Reconstruction and Analysis for Python. BMC Syst Biol 7, 74.

Ernst, J., Beg, Q.K., Kay, K.A., Balazsi, G., Oltvai, Z.N., and Bar-Joseph, Z. (2008). A semi-supervised method for predicting transcription factor-gene interactions in Escherichia coli. PLoS Comput Biol 4, e1000044.

Ernst, J., Vainas, O., Harbison, C.T., Simon, I., and Bar-Joseph, Z. (2007). Reconstructing dynamic regulatory maps. Mol Syst Biol 3, 74.

Galagan, J.E., Minch, K., Peterson, M., Lyubetskaya, A., Azizi, E., Sweet, L., Gomes, A., Rustad, T., Dolganov, G., Glotova, I., et al. (2013). The Mycobacterium tuberculosis regulatory network and hypoxia. Nature 499, 178–183.

Gardner, T.S., Cantor, C.R., and Collins, J.J. (2000). Construction of a genetic toggle switch in Escherichia coli. Nature 403, 339–342.

Geva-Zatorsky, N., Rosenfeld, N., Itzkovitz, S., Milo, R., Sigal, A., Dekel, E., Yarnitzky, T., Liron, Y., Polak, P., Lahav, G., et al. (2006). Oscillations and variability in the p53 system. Mol Syst Biol 2, 2006 0033.

Glass, L., and Kauffman, S.A. (1973). The logical analysis of continuous, non-linear biochemical control networks. J Theor Biol 39, 103–129.

Goentoro, L., Shoval, O., Kirschner, M.W., and Alon, U. (2009). The incoherent feedforward loop can provide fold-change detection in gene regulation. Mol Cell 36, 894–899.

Huang da, W., Sherman, B.T., and Lempicki, R.A. (2009). Systematic and integrative analysis of large gene lists using DAVID bioinformatics resources. Nat Protoc 4, 44–57.

Huang, S., Guo, Y.P., May, G., and Enver, T. (2007). Bifurcation dynamics in lineage-commitment in bipotent progenitor cells. Dev Biol 305, 695–713.

Kavvas, E.S., Seif, Y., Yurkovich, J.T., Norsigian, C., Poudel, S., Greenwald, W.W., Ghatak, S., Palsson, B.O., and Monk, J.M. (2018). Updated and standardized genome-scale reconstruction of Mycobacterium tuberculosis H37Rv, iEK1011, simulates flux states indicative of physiological conditions. BMC Syst Biol 12, 25.

Kempner, W. (1939). Oxygen tension and the tubercle bacillus. Am Rev Tubercul 40, 157–168.

Kholodenko, B.N. (2000). Negative feedback and ultrasensitivity can bring about oscillations in the mitogen-activated protein kinase cascades. Eur J Biochem 267, 1583–1588.

Kumar, A., Phulera, S., Rizvi, A., Sonawane, P.J., Panwar, H.S., Banerjee, S., Sahu, A., and Mande, S.C. (2019). Structural basis of hypoxic gene regulation by the Rv0081 transcription factor of Mycobacterium tuberculosis. FEBS Lett.

Kursa, M.B.R. W.R. (2010). Feature Selection with the Boruta Package. Journal of Statistical Software 36.

Langmead, B., and Salzberg, S.L. (2012). Fast gapped-read alignment with Bowtie 2. Nat Methods 9, 357–359.

Love, M.I., Huber, W., and Anders, S. (2014). Moderated estimation of fold change and dispersion for RNA-seq data with DESeq2. Genome Biol 15, 550.

Luscombe, N.M., Babu, M.M., Yu, H., Snyder, M., Teichmann, S.A., and Gerstein, M. (2004). Genomic analysis of regulatory network dynamics reveals large topological changes. Nature 431, 308–312.

Mangan, S., and Alon, U. (2003). Structure and function of the feed-forward loop network motif. Proc Natl Acad Sci U S A 100, 11980–11985.

Marcus, S.A., Sidiropoulos, S.W., Steinberg, H., and Talaat, A.M. (2016). CsoR Is Essential for Maintaining Copper Homeostasis in Mycobacterium tuberculosis. PLoS One 11, e0151816.

McMahon, M.D., Rush, J.S., and Thomas, M.G. (2012). Analyses of MbtB, MbtE, and MbtF suggest revisions to the mycobactin biosynthesis pathway in Mycobacterium tuberculosis. J Bacteriol 194, 2809–2818.

Minch, K.J., Rustad, T.R., Peterson, E.J., Winkler, J., Reiss, D.J., Ma, S., Hickey, M., Brabant, W., Morrison, B., Turkarslan, S., et al. (2015). The DNA-binding network of Mycobacterium tuberculosis. Nat Commun 6, 5829.

Murray, C.J., Ortblad, K.F., Guinovart, C., Lim, S.S., Wolock, T.M., Roberts, D.A., Dansereau, E.A., Graetz, N., Barber, R.M., Brown, J.C., et al. (2014). Global, regional, and national incidence and mortality for HIV, tuberculosis, and malaria during 1990-2013: a systematic analysis for the Global Burden of Disease Study 2013. Lancet 384, 1005–1070.

Muttucumaru, D.G., Roberts, G., Hinds, J., Stabler, R.A., and Parish, T. (2004). Gene expression profile of Mycobacterium tuberculosis in a non-replicating state. Tuberculosis (Edinb) 84, 239–246.

Nelder, J.A.M. R. (1965). A simplex method for function minimization. The Computer Journal 7.

Novak, B., and Tyson, J.J. (2008). Design principles of biochemical oscillators. Nat Rev Mol Cell Biol 9, 981–991.

Paquette, S.M.L. K.; Longabaugh, W.J.R. (2016). BioTapestry now provides a web application and improved drawing and layout tools. F1000Research 5.

Park, H.D., Guinn, K.M., Harrell, M.I., Liao, R., Voskuil, M.I., Tompa, M., Schoolnik, G.K., and Sherman, D.R. (2003). Rv3133c/dosR is a transcription factor that mediates the hypoxic response of Mycobacterium tuberculosis. Mol Microbiol 48, 833–843.

Peterson, E.J., Bailo, R., Rothchild, A.C., Arrieta-Ortiz, M.L., Kaur, A., Pan, M., Mai, D., Abidi, A.A., Cooper, C., Aderem, A., et al. (2019). Path-seq identifies an essential mycolate remodeling program for mycobacterial host adaptation. Mol Syst Biol 15, e8584.

Prosser, G., Brandenburg, J., Reiling, N., Barry, C.E., 3rd, Wilkinson, R.J., and Wilkinson, K.A. (2017). The bacillary and macrophage response to hypoxia in tuberculosis and the consequences for T cell antigen recognition. Microbes Infect 19, 177–192.

Reiss, D.J., Baliga, N.S., and Bonneau, R. (2006). Integrated biclustering of heterogeneous genome-wide datasets for the inference of global regulatory networks. BMC Bioinformatics 7, 280.

Rustad, T.R., Harrell, M.I., Liao, R., and Sherman, D.R. (2008). The enduring hypoxic response of Mycobacterium tuberculosis. PLoS One 3, e1502.

Rustad, T.R., Minch, K.J., Ma, S., Winkler, J.K., Hobbs, S., Hickey, M., Brabant, W., Turkarslan, S., Price, N.D., Baliga, N.S., et al. (2014). Mapping and manipulating the Mycobacterium tuberculosis transcriptome using a transcription factor overexpression-derived regulatory network. Genome Biol 15, 502.

Rustad, T.R., Sherrid, A.M., Minch, K.J., and Sherman, D.R. (2009). Hypoxia: a window into Mycobacterium tuberculosis latency. Cell Microbiol 11, 1151–1159.

Schnappinger, D., Ehrt, S., Voskuil, M.I., Liu, Y., Mangan, J.A., Monahan, I.M., Dolganov, G., Efron, B., Butcher, P., Nathan, C., et al. (2003). Transcriptional Adaptation of Mycobacterium tuberculosis within Macrophages: Insights into the Phagosomal Envrionment (J Exp Med).

Schulz, M.H., Devanny, W.E., Gitter, A., Zhong, S., Ernst, J., and Bar-Joseph, Z. (2012). DREM 2.0: Improved reconstruction of dynamic regulatory networks from time-series expression data. BMC Syst Biol 6, 104.

Sherman, D.R., Voskuil, M., Schnappinger, D., Liao, R., Harrell, M.I., and Schoolnik, G.K. (2001). Regulation of the Mycobacterium tuberculosis hypoxic response gene encoding alpha - crystallin. Proc Natl Acad Sci U S A 98, 7534–7539.

Shiraishi, T., Matsuyama, S., and Kitano, H. (2010). Large-scale analysis of network bistability for human cancers. PLoS Comput Biol 6, e1000851.

Smoly, I.Y., Lerman, E., Ziv-Ukelson, M., and Yeger-Lotem, E. (2017). MotifNet: a web-server for network motif analysis. Bioinformatics 33, 1907–1909.

Sun, X., Zhang, L., Jiang, J., Ng, M., Cui, Z., Mai, J., Ahn, S.K., Liu, J., Zhang, J., Liu, J., et al. (2018). Transcription factors Rv0081 and Rv3334 connect the early and the enduring hypoxic response of Mycobacterium tuberculosis. Virulence 9, 1468–1482.

Suzuki, R., and Shimodaira, H. (2006). Pvclust: an R package for assessing the uncertainty in hierarchical clustering. Bioinformatics 22, 1540–1542.

Tiwari, B.M., Kannan, N., Vemu, L., and Raghunand, T.R. (2012). The Mycobacterium tuberculosis PE proteins Rv0285 and Rv1386 modulate innate immunity and mediate bacillary survival in macrophages. PLoS One 7, e51686.

Tsai, M.C., Chakravarty, S., Zhu, G., Xu, J., Tanaka, K., Koch, C., Tufariello, J., Flynn, J., and Chan, J. (2006). Characterization of the tuberculous granuloma in murine and human lungs: cellular composition and relative tissue oxygen tension. Cell Microbiol 8, 218–232.

Vasudeva-Rao, H.M., and McDonough, K.A. (2008). Expression of the Mycobacterium tuberculosis acr-coregulated genes from the DevR (DosR) regulon is controlled by multiple levels of regulation. Infect Immun 76, 2478–2489.

Vignali, M., Armour, C.D., Chen, J., Morrison, R., Castle, J.C., Biery, M.C., Bouzek, H., Moon, W., Babak, T., Fried, M., et al. (2011). NSR-seq transcriptional profiling enables identification of a gene signature of Plasmodium falciparum parasites infecting children. J Clin Invest 121, 1119–1129.

Voskuil, M.I., Schnappinger, D., Visconti, K.C., Harrell, M.I., Dolganov, G.M., Sherman, D.R., and Schoolnik, G.K. (2003). Inhibition of respiration by nitric oxide induces a Mycobacterium tuberculosis dormancy program. J Exp Med 198, 705–713.

Wayne, L.G., and Hayes, L.G. (1996). An in vitro model for sequential study of shiftdown of Mycobacterium tuberculosis through two stages of nonreplicating persistence. Infect Immun 64, 2062–2069.

Wayne, L.G., and Sohaskey, C.D. (2001). Nonreplicating persistence of mycobacterium tuberculosis. Annu Rev Microbiol 55, 139–163.

Wernicke, S., and Rasche, F. (2006). FANMOD: a tool for fast network motif detection. Bioinformatics 22, 1152–1153.

Wu, M., Liu, L., and Chan, C. (2011). Identification of novel targets for breast cancer by exploring gene switches on a genome scale. BMC Genomics 12, 547.

Yuan, Y., Crane, D.D., Simpson, R.M., Zhu, Y.Q., Hickey, M.J., Sherman, D.R., and Barry, C.E., 3rd (1998). The 16-kDa alpha-crystallin (Acr) protein of Mycobacterium tuberculosis is required for growth in macrophages. Proc Natl Acad Sci U S A 95, 9578–9583.

